# Biophysical dissection of SOX18/NR2F2 transcriptional antagonism reveals mechanisms of venous differentiation and drug action in vascular malformation

**DOI:** 10.1101/2025.04.25.650344

**Authors:** Matthew S. Graus, Sawan Kumar Jha, Jieqiong Lou, Annegret Holm, Yew Yan Wong, Tara Davidson, Paul Coleman, Ella Sugo, Winnie Luu, Tara Karnezis, Jennifer Gamble, Geoff McCaughan, Scott Nightingale, Joyce Bischoff, Kazuaki Maruyama, Elizabeth Hinde, Kristy Red-Horse, Mathias Francois

## Abstract

Translating genomic discoveries into therapies for rare genetic disorders remains a significant challenge, particularly for variants of unknown significance (VUS) where molecular mechanisms are unclear. This is particularly relevant in vascular malformations, where venous differentiation remains poorly understood, and the role of transcription factors in specifying venous identity is only beginning to be elucidated.

Here, we combine live-cell single-molecule imaging with genomics-based approaches to uncover a biophysical mechanism of transcription factor antagonism that underpins venous identity. We show that SOX18 and NR2F2 antagonistically co-regulate venous differentiation through dynamic feedback between their nuclear populations. This interaction is disrupted in vascular malformation syndrome caused by a *de novo* heterozygous *NR2F2* mutation, presenting with aberrant vascular integrity and bleeding. Treatment with an FDA-approved drug—known to inhibit SOX18—led to marked clinical improvement in the proband.

To dissect the molecular mechanism underlying this mutation and the drug response, we used human embryonic stem cells (hESCs) engineered to carry the proband’s NR2F2 variant. These cells exhibited impaired hESC to venous differentiation with no effect on artery EC differentiation. *In silico* modelling and live-cell molecular imaging revealed that the NR2F2 variant is hyper-mobile, fails to form homodimers, and cannot recruit SOX18, thereby disrupting a critical transcriptional antagonism that underpins venous endothelial identity. We demonstrate that targeted pharmacological inhibition of SOX18 restores this regulatory balance in hESC-derived venous endothelial cells, rescuing both gene expression and NR2F2 binding dynamics at the single-molecule level. Together, this study uncovers a biophysical mechanism of transcription factor antagonism that governs venous differentiation and offers a framework for developing targeted therapies for rare vascular malformations.

## Introduction

Venous differentiation during development is a complex process governed by a network of transcription factors (TFs) that orchestrate the specification and maturation of venous endothelial cells. Despite significant advances in our understanding of arterial and venous identity, knowledge gaps persist regarding the precise regulatory mechanisms and interactions among TFs involved in venous lineage commitment. Notably, the interplay between SOX18 and NR2F2 TFs has emerged as a critical area of study, as these factors appear to play complementary roles in defining venous and lymphatic endothelium identity. Understanding this interplay may provide insights into the molecular underpinnings of vascular development and potential therapeutic targets for vascular-related diseases.

The nuclear receptor subfamily 2 group F member 2 (COUPTFII/NR2F2) is a TF required for vein development in embryonic mice, zebrafish and xenopus^1–5^. Genetic disruption approaches have demonstrated that homozygous loss of NR2F2 is embryonically lethal, and heterozygous loss is attributed to vascular anomalies^6,7^. It can also block the natural vein to artery transition required for coronary artery formation^8^. Although little is known about its function in human vein development, NR2F2 may be essential for life since it has a probability of being loss-of-function intolerant (pLI = 0.99, Genome Aggregation Database (gnomAD) v 4.1.0). Due to the inflexibility of NR2F2 variants, many are associated with or are genetic components of several human diseases, including congenital heart disease^9–11^ and congenital diaphragmatic hernia^12^, among other developmental defects^13^.

Evidence shows that NR2F2 supports venous endothelial cell (EC) differentiation though two potential mechanisms: 1. Regulating cell identity genes—promoting venous and inhibiting arterial genes^1,14,15^ and 2. enhancing cell cycle genes^8,14,15^. Deletion of one copy in coronary ECs during embryonic development drastically decreases EC numbers and proliferative capabilities^8^. Interactions with protein partners, such as GATA2 or NKX2-3, modulate part of NR2F2 activity^16,17^. In addition, the *Nr2f2* and *Sox18* genetic pathways cooperate in venous progenitor cells^18^, to initiate the early steps of lymphatic EC specification^19^ through the direct control of *Prox1* gene transactivation.

SOX18 is a member of the SOX (SRY-related HMG-box) family of TFs that is essential for different aspects of EC differentiation. It has been shown to regulate vessel calibre via genetic cross talk with VEGF-D^20^ and influences the regulation of EC contact^21,22^. During embryogenesis, SOX18 expression is essential to induce the early steps of lymphatic specification, partly through the induction of *Prox1* transcription^23^. Its expression is subsequently downregulated once the blood and lymphatic vascular networks are established. However, the interplay between NR2F2 and SOX18 TFs to coordinate and modulate their activities in veins and lymphatic ECs remains elusive.

Under pathological conditions in adults, SOX18 is reactivated and plays a role in modulating endothelial progenitor cell differentiation during solid tumour growth^24–26^. Consequently, mutations in SOX18 have been identified for their involvement in rare human vascular disorders, including hypotrichosis-lymphedema-telangiectasia-renal syndrome (HLTRS) ^27,28^ and cancer metastasis^29^. Beyond these conditions, SOX18 has also been shown to play a role in the formation of abnormal vasculature as seen in infantile hemangioma (IH), a vascular neoplasm characterized by rapidly growing, aberrant vessels. Notably, SOX18 is a key regulator of hemangioma stem cells to hemangioma endothelial cell differentiation^30,31^. This mechanism is further supported by the development of a small molecule inhibitor, SM4^18,32^ and the evidence that two FDA-approved therapies (propranolol and atenolol) are currently used to treat IH^30,31,33^, providing compelling evidence of SOX18’s involvement in pathological ECs differentiation and highlighting its potential as a therapeutic target.

Failure of these two TFs to carry out their functions often leads to lethal outcomes, due to defects that impair the cardio-vascular system and endothelial cell differentiation or function^34,35^. Here, we focus on a young proband with new *de novo* mutation in NR2F2 and the functional characterisation of this genetic insult at the molecular and cellular levels. *De novo* genetic variants have been implicated in the development of vascular malformations, with a prevalence of 1.2% to 1.5% in children^36,37^. Each disease needs to be assessed individually to identify genetic vulnerabilities and elucidate the etiological components, thereby determining treatment options. For this reason, developing assays to perform functional validation alongside next-generation sequencing technologies is essential to discovering biology associated with variants of unknown significance (VUS) and, therefore, potentially providing patients with personalized therapeutic options. There is a growing interest in VUS that alter the activity of genome effectors such as TFs, which are key molecular switches essential to proper embryonic development. TFs often operate within combinatorial networks, where opposing activities fine-tune lineage specification and gene expression. A classic example is the mutual antagonism between PU.1 and GATA1 in hematopoiesis^38,39^, where each TF represses the other’s activity to direct lineage choice. Similarly, the opposing functions of FOXO and MYC regulate the balance between cellular quiescence and proliferation^40^, while SOX2 and TCF/LEF factors exert context-dependent control over Wnt-responsive enhancers during early development^41^. These mechanisms exemplify how TFs can co-occupy shared regulatory elements yet drive divergent outcomes.

Here, we uncover an antagonistic co-regulation between SOX18 and NR2F2 essential for venous endothelial identity, by combining next-generation sequencing, *in silico* modelling, and quantitative live-cell molecular imaging. This integrated approach reveals TFs coordinated activity which disruption underlies a rare vascular syndrome. To investigate the functional consequences of a de novo heterozygous NR2F2 variant of unknown significance (VUS) identified in a proband with vascular anomalies, we employed a human embryonic stem cell (hESC) - to vascular endothelial cell (hVEC) differentiation model^42^. Notably, the proband exhibited clinical improvement following treatment with propranolol—a drug with known off-target inhibitory activity against SOX18—which led us to explore whether pharmacological disruption of SOX18 could compensate for the dysfunction of NR2F2.

Our molecular approach reveals that the VUS results in a loss-of-function NR2F2 protein that impairs venous endothelial differentiation by disrupting its interaction with SOX18, thereby uncoupling the transcriptional antagonism essential for balanced gene regulation that maintains venous identity. This mechanistic insight prompted us to test whether SOX18 inhibition could restore a transcriptional balance and phenotypic rescue. Using hESC-derived hVECs and single-molecule imaging, we demonstrate that pharmacological blockade of SOX18 functionally compensates for the NR2F2 mutation, providing proof of concept for targeted intervention via modulation of transcription factor interplay.

## Results

### NR2F2 and SOX18 influence each other’s molecular behavior

Despite pervious characterization of the genes regulated by *Sox18* and *Nr2f2* during endothelial cell development^4,43^, there is little understanding of how these two TFs biophysically and biochemically operate to instruct gene transcription of the venous or lymphatic program. Here, we investigate the molecular interactions and transcriptional co-regulation on a shared regulatory program of NR2F2 and SOX18. Analysis of a publicly available single-cell RNA sequencing (scRNA-seq) dataset of hESCs to arterial or venous ECs differentiation^42^, as well as a scRNA-seq atlas of mouse embryos at E8.5,^44^ identifies a subset of the venous population that co-expresses both NR2F2 and SOX18 (**Supp Fig. 1A-B**). We also observed co-expression in EC from vascular anomaly specimens, including infantile hemangioma (IH), venous malformations (VM), and arteriovenous malformations (AVMs) (**Supp Fig. 1C-E**). These observations prompted us to investigate how potential direct protein-protein interactions (PPI) between NR2F2 and SOX18^45^ might influence the regulation of common transcriptional targets, and how such interactions may contribute to both physiological and pathological vascular states.

**Fig 1:**
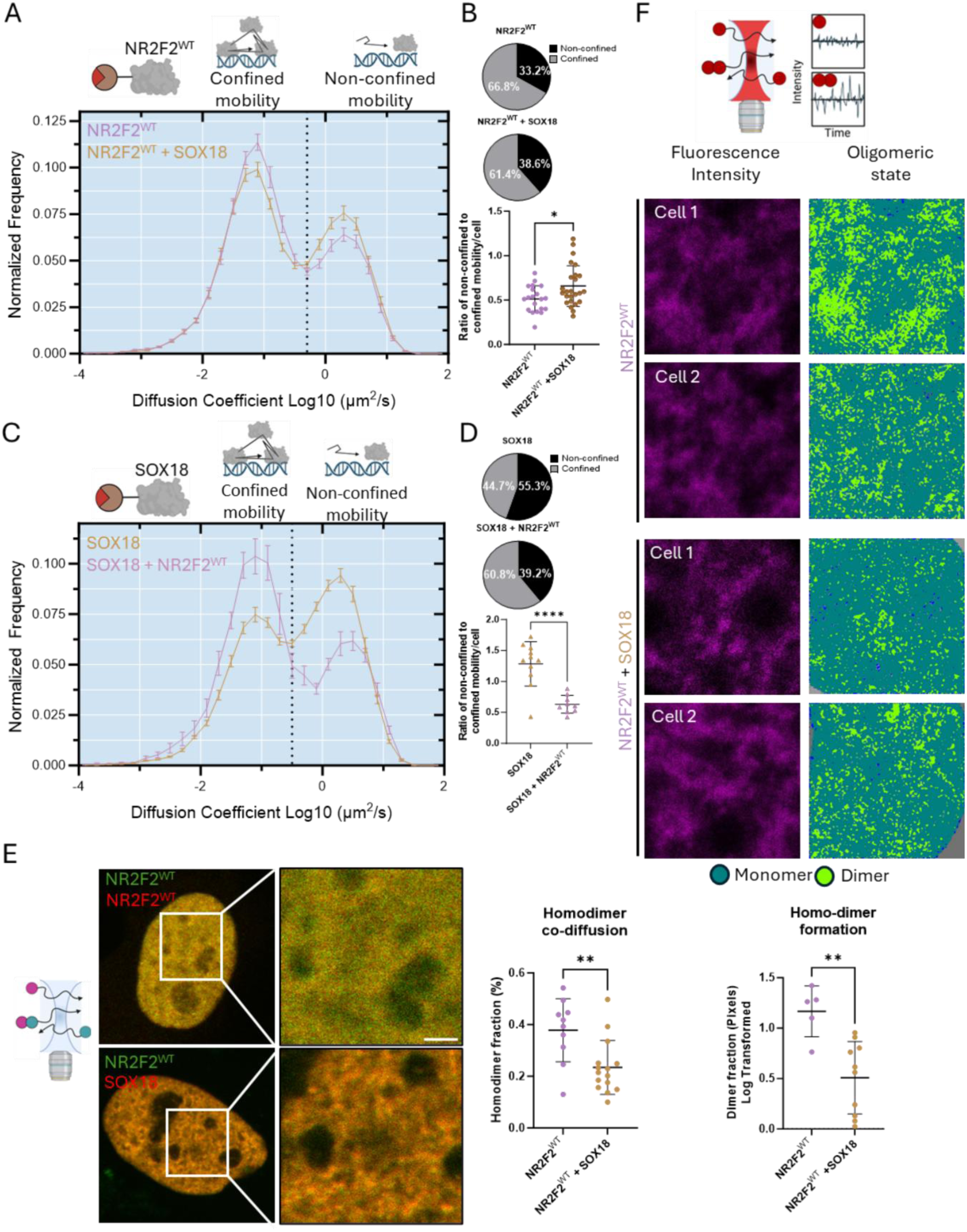
NR2F2 & SOX18 molecular interactions effect nuclear navigation. **A** Diffusion mobility graph from SMT comparing NR2F2^WT^ (purple) and NR2F2^WT^ + SOX18 (orange). **B** Pie charts represent the proportion of the population that falls into either confined mobility or non-confined mobility based on diffusion coefficient. SMT performed on NR2F2. Ratio of non-confined to confined molecules per cell from A. **C** Diffusion mobility graph from SMT comparing SOX18 (orange) and SOX18 + NR2F2^WT^ (purple). **D** Pie charts represent the proportion of the population that falls into either confined mobility or non-confined mobility based on diffusion coefficient. SMT performed on SOX18. Ratio of non-confined to confined molecules per cell from C. **E** Homo-dimer co-diffusion of NR2F2^WT^ or NR2F2^WT^ + SOX18 in the nucleus of cell. Nuclear regions of interest scale bar is 1um. **F** Homo-dimer formation measured by N&B. Fluorescent intensity of NR2F2^WT^ (top row) and corresponding oligomeric state map (bottom) for either NR2F2^WT^ or NR2F2^WT^ + SOX18. For panel E n > 11 cells and for panel F n > 5 cell, statistical significance was determined by Welch’s T-test, ** p < 0.005.

To explore a possible direct protein interaction between these two TFs, we modelled the SOX18/NR2F2 heterodimer using Alphafold 3 (**Supp Fig. 1F**, pLDDT = 51.1, pTM = 0.363, and ipTM = 0.145) ^46,47^. The predicted structure revealed a binding interface between the DNA-binding domain (DBD) of NR2F2 (amino acids 95-97) and the central transactivation domain (TAD) of SOX18 (amino acids 226-235) (**Supp Fig. 1F**). The model suggests that SOX18 might inhibit by NR2F2 and that NR2F2 might influence SOX18 PPIs with other factors since its TAD domain is used to interact with other proteins.

To functionally validate the modelled predictions, we used a combination of quantitative molecular imaging approaches to assess PPI (Number and & Brightness (N&B)), protein-chromatin interactions (Single Molecule Tracking (SMT)), and protein co-diffusion in living cells at single-molecule resolution (Cross raster image correlation spectroscopy (cRICS))^48^. cRICS measures the co-diffusion of two proteins and N&B measures the oligomeric state of the proteins, enabling the measurement of PPIs in real-time. By contrast, SMT measures the mobility and proportion of diffusing proteins, as well as their temporal occupancy^49–52^ (**Supp Fig. 2**).

**Fig 2:**
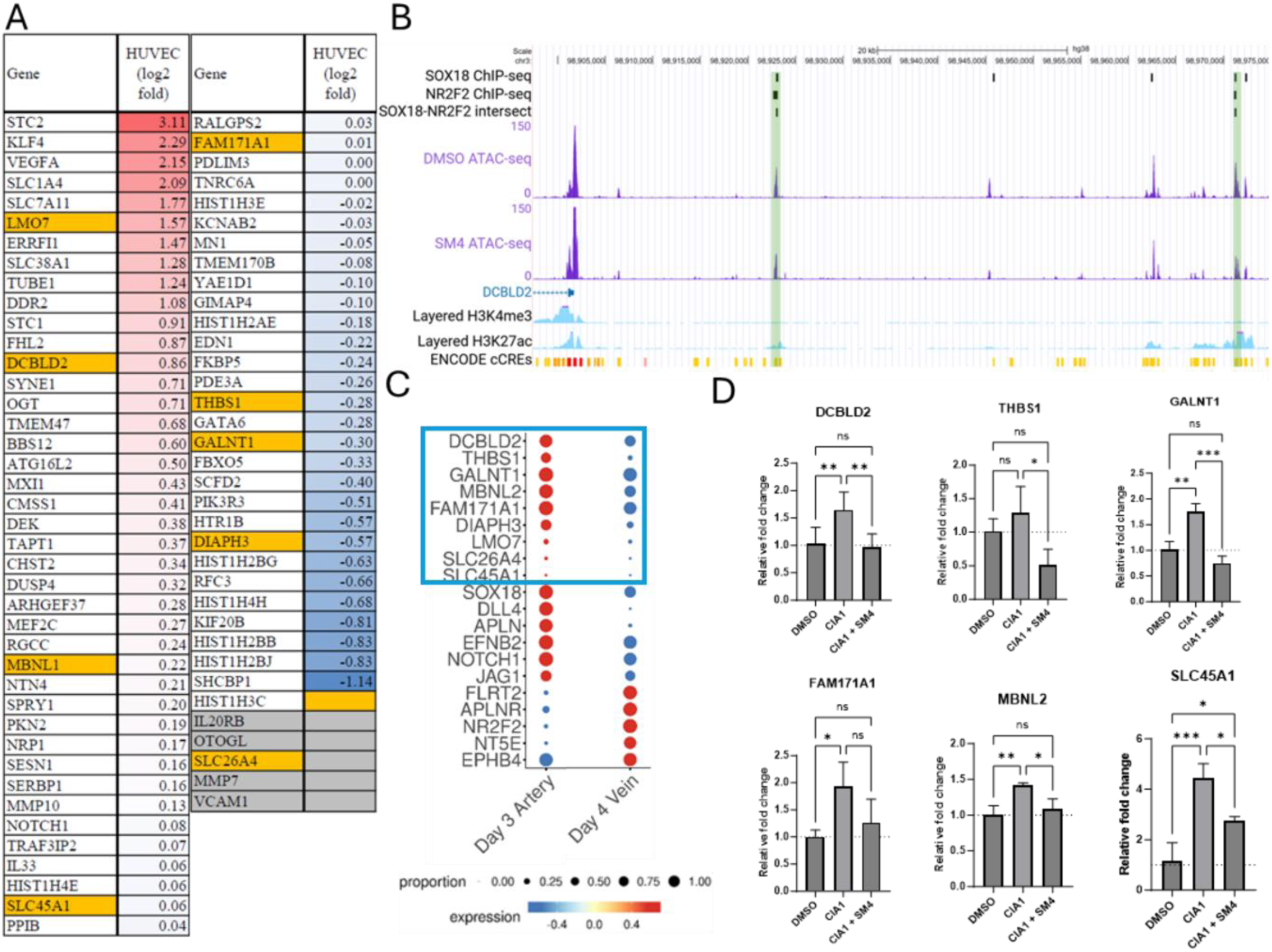
NR2F2 & SOX18 antagonistic co-regulation during venous differentiation. **A** Relative expression of a subset of genes from HUVECs treated with SM4 and measured by qPCR from Supp Table 2. Genes highlighted in yellow are the random subset of genes identified. **B** Layering of SOX18, NR2F2 ChIP-seq datasets from Sissaoui et al. (NR2F2) and Overman et al. (SOX18) (black bars), with ATAC-seq (purple), histone marks (light blue), and ENCODE cis regulatory elements map (red/orange/yellow) upstream of the DCBLD2 gene locus in HUVECs. Green boxes denote co-binding locations at distal enhancer regions by both NR2F2 and SOX18. **C** Proportion and expression of known arterio-venous identity markers. Blue box denotes a subset of genes that are found to be regulated by both NR2F2 and SOX18. Data is from Ang et al. **D** Relative gene expression of the subset of genes from panel A. HUVECs were treated with DMSO, CIA1, or CIA1 + SM4. For panel I (n = 3 repeats), statistical analysis was performed using one-way ANOVA, and significance between groups was determined using Tukey’s multiple comparison post hoc test, * p < 0.05, ** p < 0.005, *** p < 000.5, **** p < 0.0001.

This combination of assays was performed using NR2F2^WT^-halotag in the presence of SOX18 or the reciprocal condition, SOX18-halotag in the presence of NR2F2^WT^. SMT showed that when both TFs are together in the same nucleus, there is a significant change in the proportion of the TF population that interacts with the chromatin. When imaging NR2F2^WT^ in the presence of dark SOX18, the nuclear receptor exhibits more diffusive behaviour compared to when it is expressed alone (**Fig. 1A–B**). In contrast, SOX18 displays reduced mobility in the presence of dark NR2F2^WT^, compared to its behaviour in the absence of NR2F2^WT^ (**Fig. 1C-D**). These changes in mobility observed by SMT support a model in which SOX18 inhibits the ability of NR2F2 to stall on DNA while NR2F2 has the opposite effect on SOX18 stimulating its binding to DNA. Furthermore, cRICS and N&B analysis reveal a reduction in both homo-dimer formation and co-diffusion of NR2F2 homo-dimers in the presence of SOX18 (**Fig. 1E-F**). This further supports a direct interaction between the SOX18 and NR2F2 TFs and a dynamic competition model, whereby these two TFs antagonistically regulate access to shared genomic loci or chromatin territories. Collectively, these observations describe how protein-partners can influence each other’s biophysical behaviours (summarized in **Table 1**). Characterizing this interplay prompted us to explore whether these TFs may differentially modulate a shared gene regulatory program in endothelial cells.

**Table 1:** Summary of biophysical experiments and biological interpretations for NR2F2 and SOX18 interactions.

| Interplay between SOX18 and NR2F2 |  |  |  |
| --- | --- | --- | --- |
| SMT (=TF mobility and dwell time) | Observation | Biological interpretation | Fig |
| NR2F2<sup>WT</sup> non-confined mobility in presence or absence of SOX18 | NR2F2<sup>WT</sup> + SOX18: decreased proportion | NR2F2<sup>WT</sup> /SOX18 interplay leads to a more diffusive behavior of NR2F2<sup>WT</sup>, possibly via sequestration away from chromatin or competitive binding | Fig 1A-B |
| NR2F2<sup>WT</sup> confined mobility (protein-protein or protein-chromatin interaction) in presence or absence of SOX18 | NR2F2<sup>WT</sup> + SOX18: increased proportion |  |  |
| SOX18 non-confined mobility (diffusing) in presence or absence of NR2F2<sup>WT</sup> | SOX18+NR2F2<sup>WT</sup>: decreased proportion | NR2F2<sup>WT</sup> stabilizes SOX18 binding leading to increased confined fraction and less diffusive behavior possibly via increase in SOX18 homodimer formation | Fig 1C-D |
| SOX18 confined mobility (protein-protein or protein-chromatin interaction) in presence of NR2F2<sup>WT</sup> | SOX18+ NR2F2<sup>WT</sup>: increased proportion |  |  |
| N&B (=PPI, oligomeric status) and cRICS (co-diffusion) | Biophysical Observation | Biological interpretation | Fig |
| SOX18 / NR2F2<sup>WT</sup> heterodimer fraction relative to NR2F2<sup>WT</sup> only | NR2F2<sup>WT</sup> + SOX18: significantly decreased | SOX18 interferes with NR2F2<sup>WT</sup> homodimer formation | Fig 1E-F |

**Table 2:** Summary of biophysical experiments and biological interpretations of NR2F2 and SOX18 interactions.

| Interference mechanism caused by the variant |  |  |  |
| --- | --- | --- | --- |
| N&B (=PPI, oligomeric status) and cRICS (co-diffusion) | Biophysical Observation | Biological interpretation | Fig |
| NR2F2<sup>WT</sup> homodimer formation and co-diffusion compared to NR2F2<sup>Mut</sup> | NR2F2<sup>Mut</sup>: significantly decreased | c994del does not form homodimers | Fig. 5B,C |
| NR2F2<sup>WT</sup> homodimer co-diffusion compared to NR2F2<sup>Mut</sup> in the presence of SOX18 | NR2F2<sup>Mut</sup>: significantly decreased | The presence of SOX18 does not affect homodimer co-diffusion | Fig. 5F |
| NR2F2<sup>WT</sup> homodimer formation compared to NR2F2<sup>Mut</sup> in the presence of SOX18 | NR2F2<sup>Mut</sup>: significantly decreased | The presence of SOX18 does not affect homodimer formation | Fig. 5G |
| SMT (=TF mobility and dwell time) | Biophysical Observation | Biological interpretation | Fig |
| NR2F2<sup>WT</sup> non-confined mobility (diffusing) compared to NR2F2<sup>Mut</sup> | NR2F2<sup>WT</sup>: decreased proportion | NR2F2<sup>WT</sup> can interact with chromatin and other proteins whereas NR2F2<sup>Mut</sup> is unable to do so | Fig. 5D,E |
| NR2F2<sup>WT</sup> confined mobility (protein-protein or protein-chromatin interaction) in presence of NR2F2<sup>Mut</sup> | NR2F2<sup>WT</sup>: increased proportion |  |  |
| SOX18 non-confined mobility (diffusing) in presence or absence of NR2F2<sup>WT</sup> | SOX18 + NR2F2<sup>WT</sup>: decreased proportion | NR2F2<sup>WT</sup> stabilizes SOX18 binding leading to increased confined fraction and less diffusive behavior possibly via increase in SOX18 homodimer formation, but does not influence NR2F2<sup>Mut</sup> | Fig. 5H,I |
| SOX18 confined mobility (protein-protein or protein-chromatin interaction) in presence of NR2F2<sup>WT</sup> | SOX18 + NR2F2<sup>WT</sup>: increased proportion |  |  |
| SOX18 non-confined mobility (diffusing) in presence or absence of NR2F2<sup>Mut</sup> | SOX18 + NR2F2<sup>Mut</sup>: no change |  | Fig. 5H,I |
| SOX18 confined mobility (protein-protein or protein-chromatin interaction) in presence of NR2F2<sup>Mut</sup> | SOX18 + NR2F2<sup>Mut</sup>: no change |  |  |
| NR2F2<sup>Mut</sup> non-confined mobility diffusing) relative to NR2F2<sup>Mut</sup> + SOX18 | NR2F2<sup>Mut</sup> + SOX18: no change = no interaction with SOX18 | Majority of c994del molecules diffuse rather than interact = failure to efficiently select target genome location | Fig. 5J,K |
| NR2F2<sup>Mut</sup> confined mobility (protein-protein or protein-chromatin interaction) relative to NR2F2<sup>Mut</sup> + SOX18 | NR2F2<sup>Mut</sup> + SOX18: no change = no interaction with SOX18 |  |  |

### NR2F2 and SOX18 antagonistically regulate a subset of genes of the arteriovenous program

To explore a potential antagonistic modulation of genes between NR2F2 and SOX18, we integrated two previously published RNA-seq datasets, comparing known EC identity genes. The first column examines HUVECs with NR2F2 deletion (KO) and the second dataset examines HUVECs with SOX18 inhibition (treated with a small molecule inhibitor, SM4^15,18^ (**Supp Fig. 3A**). We observed that the gene response to the perturbed activity of one TF consistently leads to an opposite response when the other TF is perturbed. This antagonistic response for a subset of specific arteriovenous markers prompted us to expand our analysis to a larger gene set.

**Fig 3:**
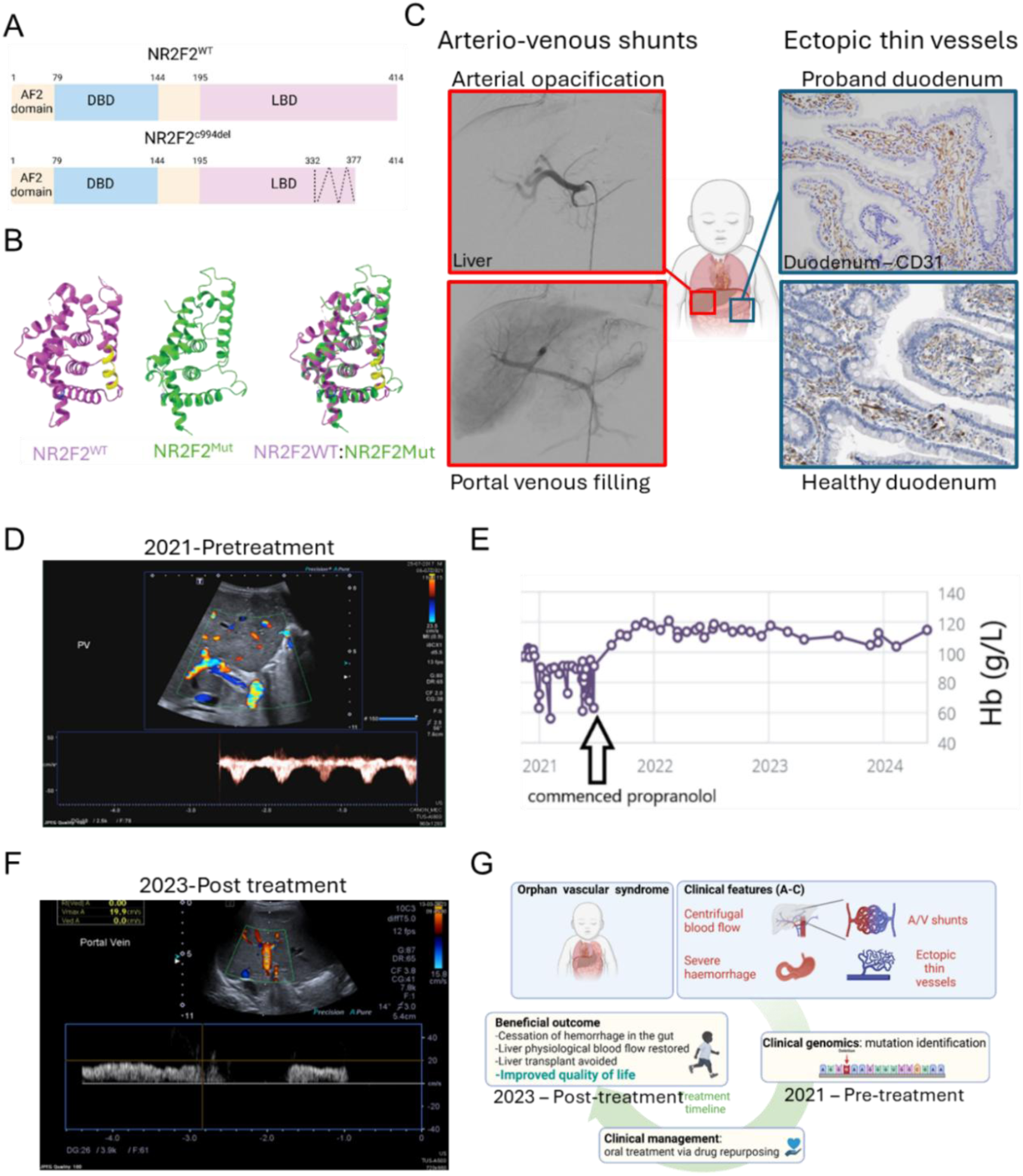
A *de novo* NR2F2 pathogenic variant causes an arterio-venous disorder that is responsive to SOX18 inhibition with propranolol. **A** Schematic representation of NR2F2^WT^ and NR2F2^Mut^. **B** Modeled structure of the LBD of NR2F2^WT^ (purple) and NR2F2^Mut^ (green). The crystal structure of the LBD was previously produced by Kruse et al. This comparison uncovers conformation difference (yellow). **C** Images from the proband’s hepatic angiography. Arterial opacification (top, red boarder) and subsequent portal venous filling (bottom, red boarder) showing diffuse arterioportal shunting. Tissue section of duodenum (H&E and counter-stained for CD31), reveals an increased number of thin-walled ectopic vessels in the proband (top, blue boarder) and healthy duodenum from a 50 year old male (bottom, blue boarder). Healthy duodenum image is the Human Atlas. **D** Doppler ultra-sound of the portal vein prior to propranolol treatment shows hepatofugal (reversed) flow in the portal vein consistent with portal hypertension. **E** Clinical tracking of hemoglobin levels prior and post propranolol treatment. **F** Doppler ultra-sound of portal vein post propranolol treatment showing restoration of physiological blood flow (hepatopetal). **G** Treatment and recovery timeline; including identification of clinical features, identification of NR2F2 variant, followed by propranolol treatment and successful pharmacological management of the condition.

To determine common target genes for these two TFs on a genome-wide scale, we combined four publicly available sequencing datasets. We layered NR2F2 and SOX18 chromatin immunoprecipitation sequencing (ChIP-seq) data sets^15,48^ with ChiP-seq datasets for marks of active gene expression (e.g. H3K27ac and H3Ke4me3 ENCODE consortium^53^) and intersected these peaks with assay for transposase-accessible chromatin (ATAC-seq) results from HUVECs that were either untreated or treated with SOX18 inhibitor, SM4^18,32^. This approach enabled us to identify regulatory regions that are co-bound by SOX18 and NR2F2 and that change their accessibility upon SOX18 inhibition. We identified 139 regions with an enhancer-like signature (ENCODE cCRE^53^ and H3K27ac, **Supp Fig. 3B**, yellow bar and light blue track) that are recruited by SOX18 and NR2F2 and less accessible upon SM4 treatment (**Supp Fig. 3B**, green bar). Peak to gene analysis^54,55^, assigned these enhancers to probable transcription start sites (TSS) and revealed 245 putative target genes (**Supp Table 1**). Cross checking known markers from that reference gene list, with a publicly available scRNA-seq dataset of hESC to endothelial differentiation^42^, reveals that both SOX18 and NR2F2 potentially co-regulate the arterio-venous program – i.e. regulate genes enriched in either artery or vein ECs (**Supp Fig. 3B**) and narrowed down the list of candidate genes to 70 genes with an arterial or venous specific expression profile (**Supp Table 2**). This observation suggests that SOX18 might alter chromatin availability in a way that disfavours NR2F2 co-binding, hence explaining the change in protein diffusion previously observed with SMT **(Fig. 1A-D)**.

To further validate SOX18-dependency of these 70 target genes, we performed qPCR analysis on HUVECs treated with SM4 (**Fig. 2A**). This approach revealed that these genes are either trans-activated or repressed by SOX18. We then assessed whether a subset of these genes is co-regulated by both NR2F2 and SOX18. A random subset of these 70 genes (**Fig. 2A**, yellow highlights) was selected for further investigation. For example, at the DCBLD2 locus, overlaying ChIP-seq and ATAC-seq datasets revealed two binding sites that correspond to both NR2F2 and SOX18 (ChIP-seq in HUVECs, black boxes). These regions both show a change in chromatin accessibility upon SM4 treatment (ATAC-seq of HUVECs treated with SM4, purple tracks) and are annotated as candidate cis-regulatory elements (cCREs, yellow boxes), indicating enhancer-like function. These results show that SOX18 and NR2F2 bind overlapping enhancer regions, and that SOX18 inhibition alters chromatin accessibility at these sites (**Fig. 2B**, green boxes).

We then compared the random gene subset to a publicly available scRNA-seq dataset to hESC-to-endothelial cell differentiation, revealing an expression pattern for genes associated with an arterial phenotype (**Fig. 2C**, blue box). This suggests that disruption of SOX18/NR2F2 co-regulation has the potential to impact EC identity by dysregulation of artery enriched genes. To test this, we performed qPCR on HUVECs treated with either DMSO as a control, or the NR2F2 inhibitor CIA1^56^ alone or in combination with SM4. A striking feature of the gene expression profiles under different treatments was a clear opposite response to either NR2F2 inhibition or the simultaneous blocking of both TFs. When cells were treated in the presence of both inhibitors, there was a consistent pharmacological rescue by SOX18 inhibitors on CIA1 dysregulated targets (**Fig. 2D**, **Supp Fig. 3C**). These observations strongly suggest that NR2F2 and SOX18 jointly regulate a subset of shared direct target genes in an antagonistic manner, which plays a role in endothelial cell identity.

This mechanistic insight raised the question of whether disrupting the cooperative antagonism of these TFs during development may underlie human vascular anomalies. A rare vascular phenotype in a paediatric proband harbouring a novel NR2F2 mutation provided an opportunity to test this hypothesis.

### A *de novo* NR2F2 variant is identified in a novel vascular anomaly

In 2020, a five-year-old proband was diagnosed with multiple vascular lesions that radically impacted development and quality of life. Owing to the rarity and severity of the disease and its resistance to treatment, whole-exome sequencing (WES) was performed. WES identified a heterozygous VUS in exon 3/3 of the NR2F2 gene (c.994del (G), p.Ala332Pro). A review of the Online Mendelian Inheritance in Man (OMIM) entry for NR2F2 identified *epicanthus inversus* (a distinctive eyelid fold abnormality) as one of its clinical features, which is a relatively uncommon condition and was also observed in this proband. This prompted us to focus our analysis further on this NR2F2 variant.

The identified mutation (NM_021005.4), Chr15 (GRCh38): g.96337371del, introduces a frame-shift beginning at amino acid position 332, resulting in an alanine-to-proline substitution leading to a scramble and truncation of the ligand binding domain (LBD) (**Fig. 3A**). The c994del variant (NR2F2^c994del^, hereafter NR2F2^Mut^) has not been reported previously, making it clinically novel. Previous NR2F2 deleterious variants have been identified and shown to cause a wide range of symptoms, ranging from cardiac to sex determination defects^9–11,13^, unlike this VUS, which seems to be involved explicitly with severe vascular defects.

To investigate the potential molecular consequences of this mutation, we used AlphaFold 3 ^46,47^ to perform *in silico* modelling and examine how the frameshift mutation affects the LBD conformation. Structural predictions of the NR2F2^WT^ LBD determined that this region is critical for its homo-dimerization^57^. In contrast, the NR2F2^Mut^ confirmation showed significant deviation, with a loss of α-helical structures following amino acid 332. This suggests that the mutation causes an abnormal folding confirmation compared to the NR2F2^WT^ LBD, starting after amino acid 332 (**Fig. 3B**, **Supp Fig 4A**).

**Fig 4:**
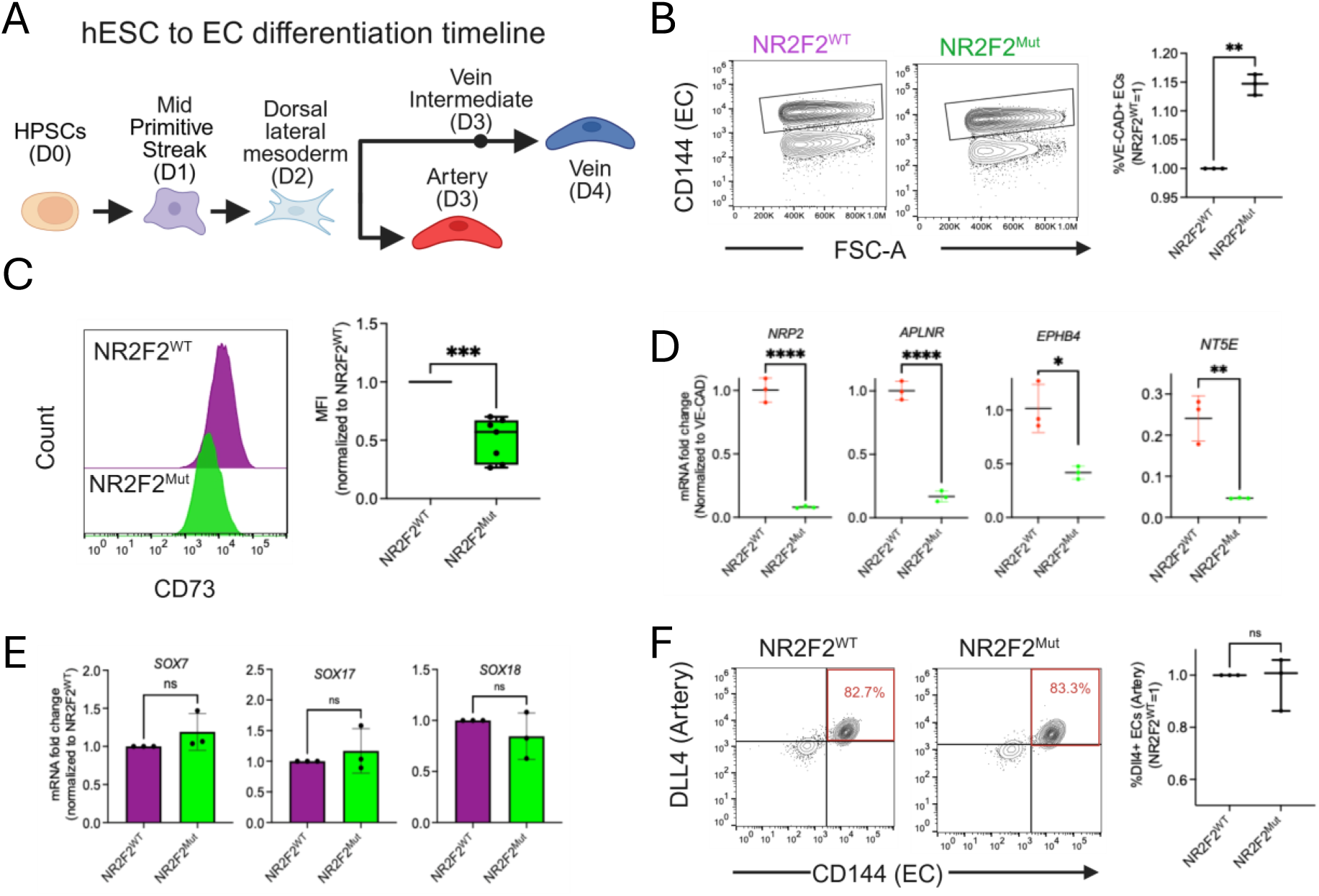
Patient’s variant impairs differentiation of hESC to venous endothelial cells. **A** Representative timeline of hESC to EC timeline. **B** Flow cytometry analysis of VE-Cadherin (pan-EC) positive ECs at day 4 of differentiation. WT (purple) Mut (green), Quantification of VE-Cadherin. **C** Histogram plots of CD73 expression in day 4 venous ECs derived from NR2F2^WT^ and NR2F2^c994del^ H1-hESC and corresponding quantification of median fluorescence intensity (MFI). **D,E** qPCR performed on day 4 venous ECs. **F** Flow cytometry analysis of artery ECs at day 4 and corresponding quantification of % DLL4+ CD144+ endothelial cells. Each data point represents independent repeats of n≥3. Statistical significance was determined by Welch’s T-test, * p < 0.05, ** p < 0.005, *** p < 0.0005, **** p < 0.0001.

WES also detected a hemizygous variant in MAP7D3 (c.1783dupA, p.Met580Asnfs*4). The variant is found in exon 10/19 of MAP7D3 gene (NM_024597.4), ChrX (GRCh38): g.136230397dup, which causes a frameshift and is expected to result in a nonsense mediated mRNA decay. This variant has been found in other individuals and is not considered a ClinVar variant on gnomAD (Genome Aggregation Database (gnomAD) v 4.1.0). The MAP7D3 gene has not been reported as a key regulator of vascular development, and there is no literature on the role of this factor in the endothelium. Therefore, we focused our investigation on the cell-autonomous role of NR2F2 dysfunction, given its well established role in venous EC identity during embryogenesis.

We started by investigating the expression profiles of NR2F2 and SOX18 in a human embryo at CS17 (gestational age: 42-44 days). During physiological vascular development, immunofluorescence staining revealed that ECs co-expressing both TFs can be found in the liver, gut, aorta, and at the junction of the cardinal vein (**Supp Fig. 4B**). Examination of the proband’s liver and duodenum (polyp) tissue samples revealed co-expression of NR2F2 and SOX18 in CD31-positive vessels (**Supp Fig. 4C-D**). Two subsets of vessels were observed: one that only expresses SOX18 (most likely arteries based on their morphological features) and the other co-expresses both NR2F2 and SOX18 (most likely veins based on the expression of NR2F2 and morphological features) **(Supp Fig. 4C-D**).

Early indications of the disease manifested as recurrent gastrointestinal haemorrhages, commencing when the patient was a newborn. Other clinical features of the disease became apparent as the proband developed. These include multiple gastric and small bowel mucosal vascular lesions characterized by an increased number of ectopic vessels and dilated capillaries, non-cirrhotic portal hypertension associated with hepatofugal flow, and diffuse hepatic arterioportal blood shunting that did not respond to surgical shunt treatments (**Fig. 3C-D**). Due to chronic gastrointestinal haemorrhages, a porta-cath was necessary for regular blood and iron transfusions, and plans were underway to list for a liver transplantation by pre-school age.

Mutations in key developmental TFs specific to EC development have been reported to cause rare vascular diseases^60,61^. For instance, mutations in the transactivation domain of SOX18 TF are known causes of HLTRS, which has a dominant-negative effect and suppresses the endogenous function of the wild-type protein^48,62,63^. Remarkably, the FDA-approved drug propranolol has shown considerable efficacy in enhancing the clinical management of the HLTRS syndrome^31^. Since the early 2010s, this drug has been primarily used as the first-line treatment for IH, the most common vascular tumor, whose mechanism of action also relies on SOX18 inhibitory activity^30,31,33^. Given propranolol’s established safety for use in paediatric vascular disorders, the identification of its recent mode of action via SOX18 blockade, and the worsening condition of the proband with limited therapeutic options, a decision was made to use propranolol to manage the proband’s condition.

### Treatment and recovery

Propranolol treatment was initiated at a dose of 1.5mg/ml/kg in August 2021. By December 2021, haemorrhaging had resolved, and haemoglobin levels stabilized, resulting in the patient no longer requiring transfusions (**Fig. 3E**). In early 2023, a Doppler ultrasound was performed, and it determined that hepatopetal flow had been restored (**Fig. 3F**). Due to the proband no longer requiring transfusions the porta-cath was removed, and the liver transplant was cancelled. At the time of writing this manuscript the treatment has been a success, with the probands’ quality of life drastically improving, allowing for a return to a more normal and fulfilling life (**Fig. 3G**). Following the successful treatment, we aimed to unravel the underlying cellular and molecular mechanisms of propranolol’s therapeutic effect. Importantly, the co-expression of SOX18 and NR2F2 in the proband’s endothelium **(Supp Fig. 4C-D**) raises the question whether NR2F2 mutation would directly affect the interplay with SOX18 and impact their co-regulation process and whether propranolol treatment has the potential to restore this imbalanced relationship.

### NR2F2^Mut^ impairs venous but not arterial differentiation

We started by investigating the pathogenicity of the NR2F2 variant. Given its importance for vein identity in model organisms, we tested the impact of the variant on vein EC differentiation in a highly efficient human stem cell to venous and arterial EC differentiation model^42^ (**Fig. 4A**). We generated H1-hESC lines containing the proband-specific variant in a homozygous fashion (**Fig. 4A**, **Supp Fig. 5A-D**). Western blotting confirmed that the NR2F2^Mut^ lines had NR2F2 proteins at comparable expression to wild type, but, as expected, at a lower molecular weight (predicted NR2F2^WT^ = 45.5kDa (414AA) and NR2F2^c994del^ = 41.03kDa (377 AA), consistent with a premature stop codon. (**Supp Fig. 5C**).

**Fig 5:**
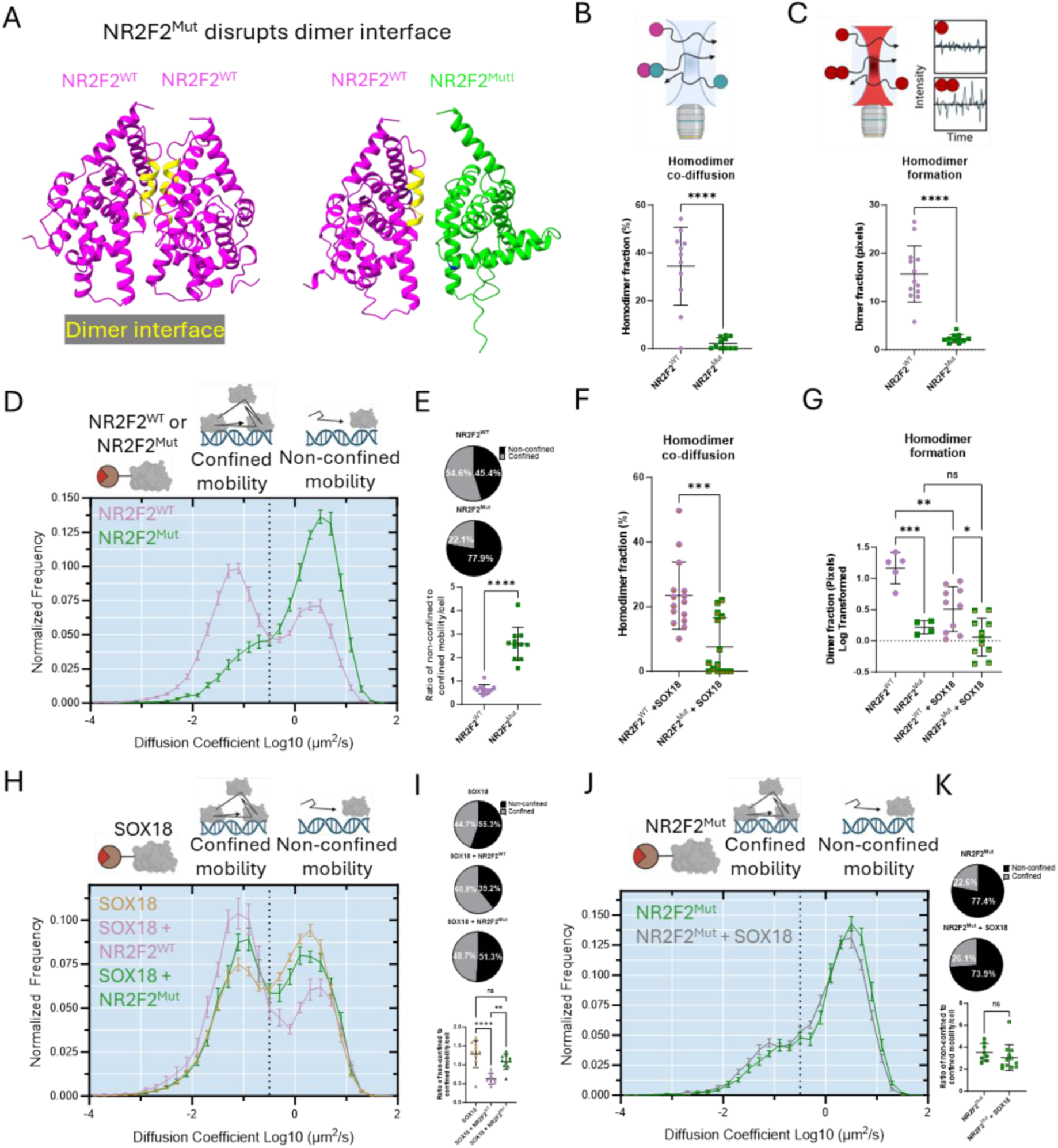
NR2F2^Mut^ variant shows altered molecular behavior and unbalances SOX18 activity. **A** Modeled dimerization of the LBD of NR2F2^WT^ (purple) and NR2F2^c994del^ (green), dimerization domains (yellow). NR2F2^WT^ or NR2F2^Mut^ measured by cRICS to assess homo-dimer co-diffusion. **B, C** NR2F2^WT^ or NR2F2^Mut^ measured by cRICS (B) and N&B (C) to assess homo-dimer diffusion and formation. **D** Diffusion mobility graph from SMT comparing NR2F2^WT^ and NR2F2^Mut^. **E** Pie charts represent the proportion of the population that falls into either confined mobility or non-confined mobility based on diffusion coefficient. Ratio of non-confined to confined molecules per cell from D. **F, G** Hetero-dimer co-diffusion of NR2F2^WT^ + SOX18 or NR2F2^Mut^ + SOX18 measured by cRICS (F) and N&B (G). **H** Diffusion mobility graph from SMT comparing SOX18 (orange), SOX18 + NR2F2^WT^ (purple), and SOX18 + NR2F2^Mut^ (green). **I** Pie charts represent the proportion of the population that falls into either confined mobility or non-confined mobility based on diffusion coefficient. SMT performed on SOX18. Ratio of non-confined to confined molecules per cell from H. **J** Diffusion mobility graph from SMT comparing NR2F2^Mut^ (green) and NR2F2^c994del^ + SOX18 (grey). **K** Pie charts represent the proportion of the population that falls into either confined mobility or non-confined mobility based on diffusion coefficient. SMT performed on NR2F2. Ratio of non-confined to confined molecules per cell from J. For panels B, C, E, F, and K; n > 12 cells, statistical analysis was performed using Welch’s T-test, * p < 0.05, ** p < 0.005, *** p < 000.5, **** p < 0.0001. For panels G and I n > 5 cells, statistical analysis was performed using one-way ANOVA, and significance between groups was determined using Tukey’s multiple comparison post hoc test, * p < 0.05, ** p < 0.005, *** p < 000.5, **** p < 0.0001.

To assess the phenotypic impact of the NR2F2 variant and its cell autonomous role, we differentiated wild type control and NR2F2^Mut^ hESCs into vein and artery ECs. The differentiation assay utilized FACS and qPCR readouts to assess cellular and molecular levels of arterial and venous differentiation markers (vein=CD73, *APLNR*, *NRP2,* and SOXFs). In line with its known role as a hallmark of venous identity, NR2F2^Mut^ caused an increase in the overall number of ECs (**Fig. 4B**) and resulted in a significantly decreased levels of the hESC-derived vein marker CD73 (**Fig. 4C**, CD73+ ECs, **Supp Fig. 5D**). The additional venous markers, *APLNR, EPHB4, NRP2*, and *NT5E* (*CD73*) were also down as measured by qPCR, further expanding FACS observations and indicating a potential defect in achieving full venous identity (**Fig. 4D**). NR2F2 deficiency also did not change the expression of SOX18 or other SOXF’s (**Fig. 4E)**. The lack of change in the SOX7 and SOX18 expression in the NR2F2 variant clones suggested that this TF is not directly upstream of SOXF members in the model of venous differentiation (**Fig 4E**). In contrast to its effect on vein endothelial cells, NR2F2^Mut^ did not influence the arterial differentiation process (**Fig. 4F**). Of interest is the finding that NR2F2^Mut^ cells make more ECs because it aligns with the patient duodenum tissue sample showing an increased number of ectopic, thin-walled vessels (**Fig. 3C**). These findings show that the NR2F2^Mut^ can autonomously interfere with EC subtype transcriptional profiles, specifically within the venous compartment. Based on the previous interplay between SOX18 and NR2F2 under physiological conditions, we next set out to determine the mode of action of the proband’s mutation.

### Molecular mechanism of NR2F2^Mut^ impaired function

Because the proband is heterozygous and has both wild type and variant alleles expressed, we modelled the effect of this abnormal configuration on homodimer formation. Crystal structure studies of the LBD of NR2F2^WT^ further confirmed the dimerization residues involve several α-helices, including 7 (amino acids 297-301), 8 (amino acid 378), 9 (amino acids 335 and 339), and 10 (amino acids 358-372), along with the loop between α-helices 8 and 9 (amino acids 323 and 324)^57^. Because the location of the frame-shift in the variant occurs at amino acid 332, these dimer residues are lost, potentially resulting in an abnormal folding conformation at the C-terminus. As expected, the *in silico* modelling indicates that the loss of dimer residues results in the inability of the variant and WT to form a dimer. The mutation results in a disordered conformation, which likely creates an unstable interface compared to NR2F2^WT^ (**Fig. 5A**, **Supp Fig 6A-C**), which could result in a global destabilization of the NR2F2 homodimer pool.

**Fig 6:**
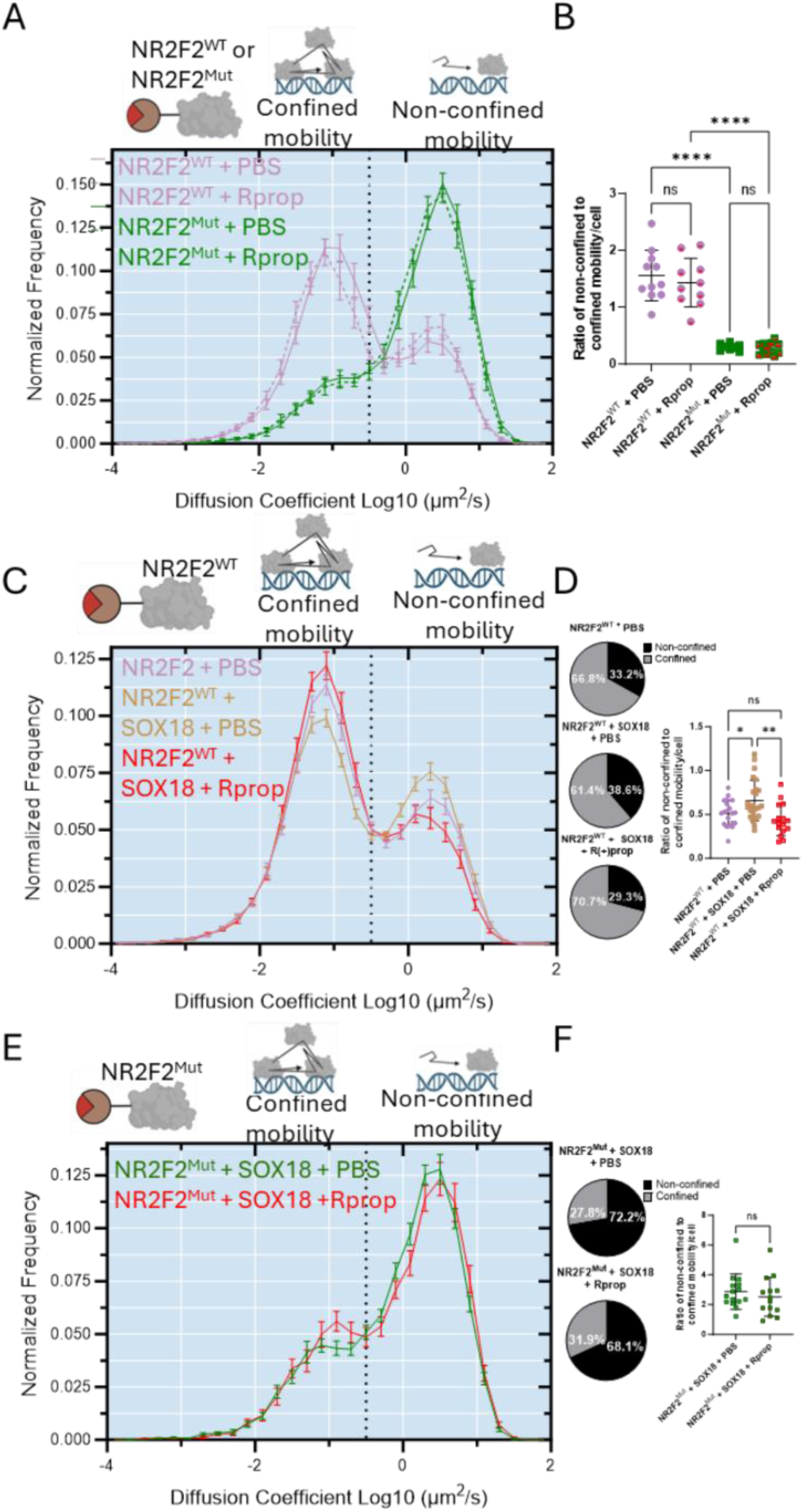
NR2F2^Mut^ molecular rescue with R(+)propranolol treatment is SOX18-dependent. **A** Diffusion mobility graph from SMT comparing NR2F2^WT^ + PBS (purple), NR2F2^WT^ + Rprop (purple, hashed), NR2F2^Mut^ + PBS (green), and NR2F2^Mut^ + Rprop (green, hashed). SMT performed on NR2F2. **B** Ratio of non-confined to confined molecules per cell from A. **C** SMT performed on NR2F2. Diffusion mobility graph from SMT comparing NR2F2^WT^ + PBS (purple), NR2F2^WT^ + SOX18 + PBS (orange), and NR2F2^WT^ + SOX18 + R(+)prop (red). **D** Pie charts represent the proportion of the population that falls into either confined mobility or non-confined mobility based on diffusion coefficient. Ratio of non-confined to confined molecules per cell from C. **E** Diffusion mobility graph from SMT comparing NR2F2^c994del^ + SOX18 + PBS (grey), and NR2F2^c994del^ + SOX18 + Rprop (red). **F** Pie charts represent the proportion of the population that falls into either confined mobility or non-confined mobility based on diffusion coefficient. SMT performed on NR2F2. Ratio of non-confined to confined molecules per cell from E. For panels B and D; n > 8 cells, for panel F; n > 14 cells, statistical significance was determined by Welch’s T-test. Statistical analysis was performed using one-way ANOVA, and significance between groups was determined using Tukey’s multiple comparison post hoc test, * p < 0.05, ** p < 0.005, **** p < 0.0001.

Consistent with the observed model, cRICS and N&B imaging showed the NR2F2^Mut^ almost completely lost its ability to co-diffuse and form homo-dimers (**Fig. 5B-C**). Following on, we next investigated how the loss of homo-dimerization affects the variant’s ability to navigate and interact within the nuclear environment. SMT imaging determined that most of the variant population has an increased diffusion profile compared to its WT counterpart (**Fig. 5D-E** (HeLa), **Supp Fig. 6D-E** (HUVEC)). Taken together, the data indicate that the variant is unable to homo-dimerize, allowing the vast majority of NR2F2^Mut^ molecules to diffuse as monomers, hence dramatically impacting the total pool of NR2F2 available to perform its function.

We next assessed whether homo-dimerization is necessary for NR2F2 to bind and interact with its environment. Temporal occupancy SMT established that the variant spends less time searching for binding sites as indicated by duration of short dwell times (**Supp Fig. 6F**), but a similar amount of time interacting when an interaction occurs as indicated by duration of longer dwell times (**Supp Fig. 6G**). Additionally, the variant has a much smaller proportion of the population bound at any time indicated by the reduction in the ratio of long to short dwell time events (**Supp Fig. 6H**). These results demonstrate that the variant causes interactions to be less stable through the loss of target gene search pattern and fewer of the population undergoing longer specific interactions with the chromatin (summary of the variant biophysical features in **Table 2**).

The biophysical characterization of the defects establishes that the NR2F2^Mut^ severely disables NR2F2 activity and interferes with its physiological and molecular function, leading us to define the variant as a loss-of-function variant. We next questioned whether the variant also impacts NR2F2’s capacity to hetero-dimerize with a known protein partner. Previous studies have shown direct interaction of NR2F2 and SOX18 in cell-based assays^18,48^, and it is known that these two TFs together induce *Prox1* expression in the venous endothelium to specify lymphatic endothelial cells^19^. However, this is a synergistic relationship, and our data suggests an antagonistic interaction with respect to gene induction **(Fig. 2D)**

To determine if NR2F2^Mut^ also lost its ability to interact with SOX18 we performed cRICS, N&B, and SMT on either NR2F2^WT^ or NR2F2^Mut^-halotag in the presence of SOX18 or the reciprocal condition. cRICS and N&B analysis of NR2F2, discussed above, identified a loss of co-diffusion and homo-dimer formation for NR2F2^Mut^ (**Fig. 5B-C**). The same trend was also observed for NR2F2^Mut^ in the presence of SOX18 (**Fig. 5F-G**). To understand how hetero-dimerization affects the diffusion profile of both

TFs, we turned to SMT. This approach revealed when NR2F2^WT^ is present SOX18 mobility becomes more confined, whereas when NR2F2^Mut^ is present there is a negligible change in SOX18 mobility (**Fig. 5H-I**). By contrast, NR2F2^Mut^ mobility remains unchanged regardless of SOX18 presence (**Fig. 5J-K**). Taken together, the observations suggest NR2F2^Mut^ most likely influences SOX18 mobility indirectly through the interference with its wild-type counterpart (summary of NR2F2/SOX18 interplay in **Table 2**).

Collectively, these findings reveal that NR2F2^Mut^ is a severe loss-of-function mutation that disrupts both homo-and hetero-dimer formation, leading to a deficiency of NR2F2. The frameshift mutation results in a disordered and destabilized LBD, which depletes the homo-dimer pool. Importantly, NR2F2^Mut^ also fails to interact productively with SOX18, suggesting that the variant may interfere with other key regulators of venous identity, akin to the mechanism reported for the SOX18 dominant-negative mutation^48^. These mechanistic disruptions likely underlie the endothelial phenotype observed in the proband, supporting a model in which balanced NR2F2-SOX18 interactions are essential for endothelial transcriptional programs and venous homeostasis. These findings prompted us to question whether the proband’s recovery following propranolol treatment may have occurred, at least in part, through a rescue mechanism that involves the pharmacological blockade of SOX18 activity.

### Drug repurposing and molecular mechanisms of treatment

Recent findings have demonstrated that both propranolol and a small molecule inhibitor of SOX18, SM4 inhibit the differentiation of hemangioma stem cells into hemangioma endothelial cells in a SOX18-dependent manner^30,31,33,64^ and also blocks SOX18 pioneering function during virus-induced LEC reprogramming in Kaposi’s sarcoma^65^. Specifically, propranolol is composed of a R(+) and S(-) enantiomer mixture at a 1:1 equimolar ratio, with the R(+) enantiomer (R(+)propranolol) devoid of beta-adrenergic effects.

Previous research has demonstrated that the R(+) enantiomer is capable of disrupting SOX18 activity ^30,31,66,67^. Given that NR2F2 and SOX18 are co-expressed in a subset of the venous endothelium and antagonistically co-regulate a common set of arterial-related genes, we sought to determine whether R(+)propranolol-induced SOX18 inhibition has the potential to rescue NR2F2 loss-of-function caused by the variant NR2F2^Mut^.

Taking advantage of molecular imaging, we first evaluated if R(+)propranolol directly engages with NR2F2 by measuring its molecular mobility and temporal interaction times using SMT and N&B. Collectively, these experiments indicate that R(+)propranolol (Rprop) does not interact with or effect NR2F2 protein since it does not disrupt a broad range of key biophysical behaviours (**Fig 6A-B, Supp Fig 7 A-D**). Because R(+)propranolol does not disrupt NR2F2, we hypothesized that the treatment of the proband would have to be through the inhibition of SOX18. To test this hypothesis, we next examined the effects of this compound on NR2F2-SOX18 interactions using SMT, N&B, and cRICS. In the absence of the drug, SMT imaging revealed that the presence of SOX18 increases the proportion of diffusing NR2F2^WT^ molecules compared to NR2F2^WT^ alone, suggesting that SOX18 is directly or indirectly disfavoring NR2F2 interaction with the chromatin. Additionally, when SOX18 activity is disrupted by R(+)propranolol, we observed a decrease in the proportion of diffusing NR2F2^WT^ molecules, indicating that interference of R(+)propranolol with SOX18 restores the chromatin-bound fraction of NR2F2^WT^ to its baseline state (**Fig. 6C-D**). Conversely, there is little change in NR2F2^Mut^ diffusion profile in the presence of SOX18 with or without R(+)propranolol (**Fig. 6E-F**). This suggests that the protective effect of R(+)propranolol is mediated at a molecular level, specifically by indirectly modulating the availability of the NR2F2^WT^ pool through an interference with SOX18 activity. In other words, this work builds on previous findings that inhibition of SOX18 by pharmacological blockade reduces NR2F2 and SOX18 interaction^32^ and establishes that SOX18 inhibition favours an increase of NR2F2^WT^ activity on the chromatin (summary of drug mode of action, **Table 3**). Having elucidated that R(+)propranolol-induced SOX18 inhibition restores a vast proportion of the NR2F2^WT^ behaviour, we next investigated whether R(+)propranolol is able to rescue the venous phenotype caused by the NR2F2 variant in a human pre-clinical model of stem cell to venous endothelial cell differentiation.

**Fig 7:**
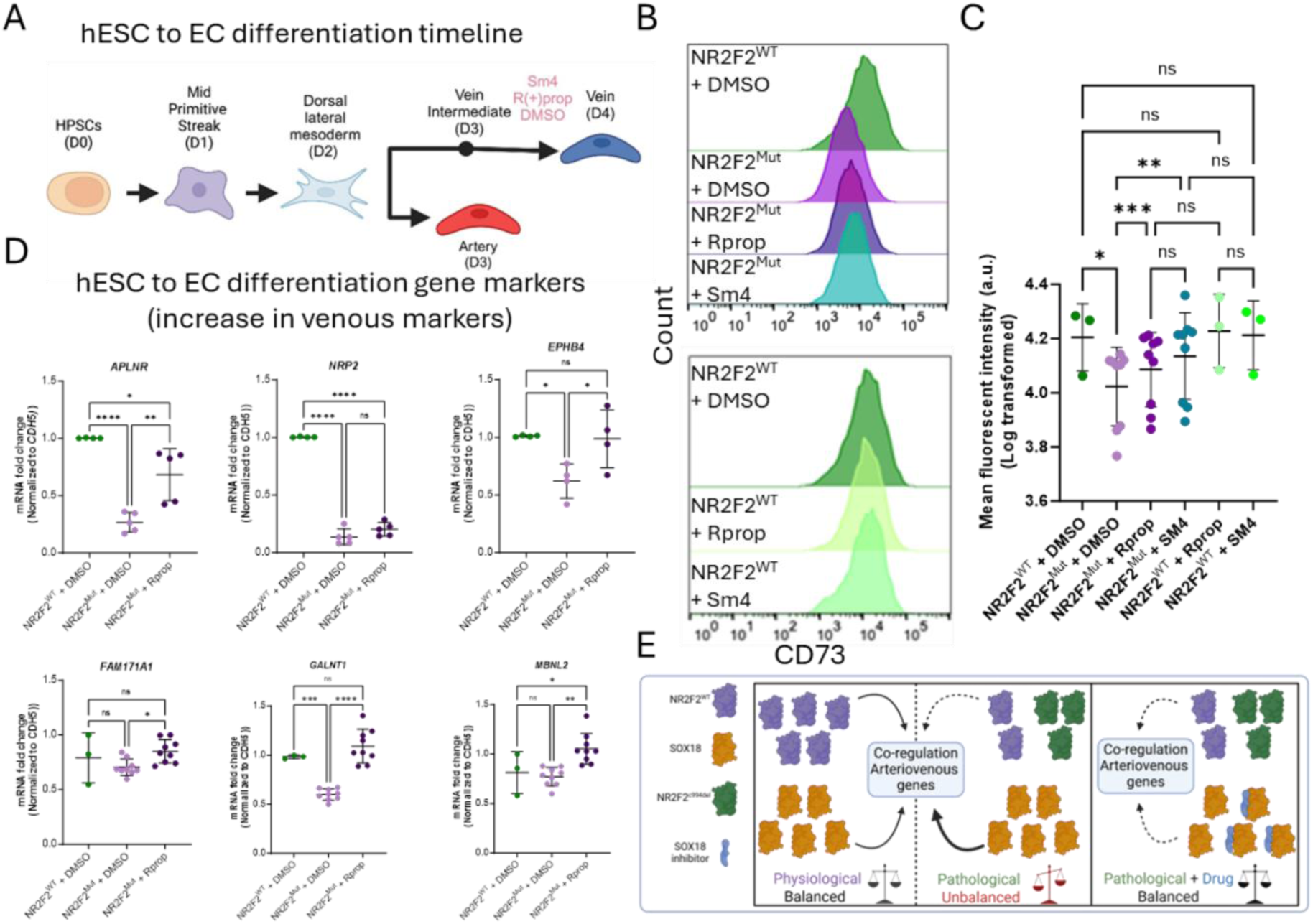
Both SOX18 small molecule inhibitor and Propranolol rescue NR2F2^Mut^-induced venous differentiation defects. **A** Schematic of hESC to EC differentiation timeline with drug treatment. **B** Histogram plots of venous derived populations and corresponding quantification of MFI. An increase of the CD73 expression occurs in the presence of SOX18 blockers despite the NR2F2 mutation. **C** Comparison of CD73 expression. The three clones carrying the mutant version of NR2F2 are pooled. **D** Relative gene expression measured by qPCR from day four venous cells. **E** Model of the proposed molecular mechanism underlying the NR2F2^c994del^ rescue. Under physiological conditions in venous cells, NR2F2 and SOX18 are co-expressed at a specific ratio. This ratio allows for an antagonistic co-regulation of arteriovenous markers in vein ECs that is essential to maintain venous identity. In the proband’s scenario NR2F2 heterozygous mutation alters the proper ratio between NR2F2^WT^ and SOX18 that then becomes unbalanced in favor of SOX18 activity, leading to an upregulation of arterial markers and a perturbation of venous identity. Treatment with a SOX18 inhibitor enables the restoration of balance between SOX18 and NR2F2 activity, allowing a smaller proportion of both TF populations to function properly at physiological levels. Panel C (n > 3 repeats of 10000 cell/cell line), statistical analysis was performed using a mixed-effects analysis, with the Geisser-Greenhouse correction, and Tukey’s multiple comparison test, * p < 0.03, ** p < 0.002, *** p < 0.0002. Panel D (n > 3 repeats/cell line), statistical analysis was performed using one-way ANOVA, and significance between groups was determined using Tukey’s multiple comparison post hoc test, * p < 0.05, ** p < 0.005, *** p < 0.005, **** p < 0.0001.

**Table 3:** Summary of biophysical experiments and biological interpretations investigating drugs effect on NR2F2 and SOX18.

| Drug mode of action |  |  |  |
| --- | --- | --- | --- |
| N&B (=PPI, oligomeric status) and cRICS (co-diffusion) | Biophysical Observation | Biological interpretation | Fig |
| NR2F2<sup>WT</sup> + PBS homodimer fraction compared to NR2F2<sup>WT</sup> + R-propranolol | NR2F2<sup>WT</sup> + R(+)propranolol: no change | R-propranolol does not affect NR2F2<sup>WT</sup> dimer formation | Supp Fig. 7A |
| NR2F2<sup>WT</sup> homodimer fraction relative to NR2F2<sup>Mut</sup> + R-propranolol | NR2F2<sup>Mut</sup> + R(+)propranolol: no change | R-propranolol does not affect NR2F2<sup>Mut</sup> oligomeric status | Supp Fig. 7A |
| SMT (=TF mobility and dwell time) | Biophysical Observation | Biological interruption | Fig |
| non-confined mobility (diffusing) of NR2F2<sup>WT</sup> or NR2F2<sup>Mut</sup> in presence or absence of R(+)propranolol | no change caused by drug treatment | R(+)propranolol does not affect NR2F2<sup>WT</sup> or variant mobility | Fig. 6A,B |
| confined mobility (diffusing) of NR2F2<sup>WT</sup> or NR2F2<sup>Mut</sup> in presence or absence of R(+)propranolol | no change caused by drug treatment |  |  |
| NR2F2<sup>WT</sup> non-confined mobility (diffusing) in presence or absence of SOX18 with or without R(+)propranolol | Adding SOX18 leads to increased NR2F2 diffusion, effects that are counteracted by R(+)propranolol | R(+)propranolol rescues NR2F2<sup>WT</sup> changes of behavior that are caused by SOX18 | Fig. 6C,D |
| NR2F2<sup>WT</sup> confined mobility (diffusing) in presence or absence of SOX18 with or without R(+)propranolol | Adding SOX18 leads to decreased NR2F2 bound fraction, effects that are counteracted by R(+)propranolol |  |  |
| NR2F2<sup>Mut</sup> non-confined mobility (diffusing) in presence of SOX18 with or without R(+)propranolol treatment | NR2F2<sup>Mut</sup> + SOX18 +/- R(+)propranolol: no change | Inhibition of SOX18 does not impact variant mobility | Fig. 6E,F |
| NR2F2<sup>Mut</sup> confined mobility (protein-protein or protein-chromatin interaction) in presence of SOX18 with or without R(+)propranolol treatment | NR2F2<sup>Mut</sup> + SOX18 +/- R(+)propranolol: no change |  |  |

### R(+)propranolol effects are endothelial cell autonomous and rescues venous defects caused by NR2F2^Mut^

To perform an orthogonal validation of data obtained from single-molecule imaging, we next employed the hESC to venous EC differentiation assay. The experimental design relies on a differentiation protocol in the presence or absence of SOX18 blockers (either SM4 or R(+)propranolol) when cells transition from vein intermediates (D3) to vein (D4). (**Fig. 7A**). To assess whether the SOX18 inhibitor exert an effect on NR2F2^Mut^-induced venous defects, we analyzed the three independent clones with the proband mutation (**Supp Fig. 5D**). These clones were evaluated for CD73 expression by FACs and venous marker expression by qPCR (**Fig. 7B-D**). Treatment with SOX18 inhibitors significantly restored CD73 expression in NR2F2^Mut^ cells relative to DMSO-treated cells while having no effect on physiological CD73 expression in NR2F2^WT^ (**Fig. 7B**, **C**). Similarly, expression of classic venous markers *APLNR, NRP2, and EPHB4*, along with three additional markers previously identified (*FAM171A, GALNT1,* and *MBNL2*) was also rescued following R(+)propranolol treatment (**Fig. 7D**). Molecular rescue was consistent for all clones tested with no significant variability in response to the inhibitors (**Supp Fig. 7E-G**) These results established that the recovery of the venous differentiation defects caused by NR2F2 variant is meditated through SOX18 blockade. The validation with this cell-based assay determined that the R(+)propranolol molecular mechanism specifically acts on the venous endothelium in a cell autonomous manned and suggests that SOX18 inhibition in the proband has contributed to a restored physiological function of the venous vasculature.

## Discussion

### The NR2F2-SOX18 axis as a regulatory hub in endothelial identity

This study centers on the hypothesis that NR2F2 and SOX18 function cooperatively to regulate venous identity and the fate of endothelial cells. Here, we show that during physiological venous differentiation, SOX18 and NR2F2 establish a regulatory equilibrium, dynamically influencing each other’s chromatin binding and transcriptional activity. Previous genetic studies have shown that these synergistic pathways regulate PROX1 and lead to the development of lymphatic endothelial cells, however the dynamics of this coordination at a biophysical and biochemical levels have remained uncharacterized. The discovery of a *de novo* NR2F2 mutation in a proband that results in severe vascular defects presented a unique opportunity to examine what happens when the equilibrium between NR2F2 and SOX18 becomes unbalanced. The proband’s improvement after the start of propranolol treatment, including cessation of hemorrhaging and the return of hepatofugal flow in the portal vein, strongly supports a model in which SOX18 inhibition compensates for NR2F2 dysfunction. *In silico* and live-cell imaging revealed that the NR2F2^Mut^ protein is unable to form homo-dimers and cannot stably interact with SOX18, resulting in the disruption of gene regulation. The loss of NR2F2 and SOX18 interactions was found to be a key driver of the proband’s vascular pathology, implicating the loss of NR2F2-SOX18 coordination as a significant contributor to this disease. Given the antagonistic transcriptional relationship between NR2F2 and SOX18, we hypothesized that inhibiting SOX18 might rebalance transcriptional regulation.

### An updated molecular mechanism for investigating TFs

Based on this study, we propose a molecular model explaining how Propranolol treatment benefited the proband. Under normal circumstances, a physiological balance between NR2F2 and SOX18 ensures the coordinated transcriptional regulation of common target genes central to arteriovenous identity. However, in the proband’s case, a heterozygous variant in NR2F2 causes only half of the pool of NR2F2 molecules to function properly, leading to a shift in balance in favour of SOX18 activity. This results in transcriptional dysregulation being over-dominated by SOX18. To correct this imbalance, a molecular strategy based on SOX18 inhibition is used to compensate for the larger pool of SOX18 compared to functional NR2F2 and to restore a relative equilibrium between the remaining functional NR2F2 and less active SOX18 molecules (**Fig. 7E**). Our findings introduce a novel concept to correct loss-of-function mutations of transcriptional regulators by targeting TFs interplay, ultimately normalizing co-regulated genes to a physiological level.

### Using molecular imaging to investigate pharmacological on-target engagement

Targeting TFs to modulate gene expression has been studied for decades. Historically, TFs have been considered “undruggable” due to their intrinsic disordered domains, lack of a traditional drug binding pocket, and dynamic nature^68^. This study highlights the potential of using quantitative molecular imaging as a strategy to perform on-target engagement. Using SMT has unlocked the ability to assess a molecular target in the presence of its inhibitor^69,70^. Here, we firmly establish that the R(+)propranolol mode of interference is mediated via SOX18 disruption, whereas NR2F2 behavior remains unaffected by this drug. Molecular imaging thus provides a powerful tool to evaluate the interaction between drugs and molecular targets, offering a direct method to validate therapies aimed at specific pathways.

### A new therapeutic pathway for congenital vascular malformations

More broadly, vascular anomalies have been linked to alterations in signaling pathways associated with the development of AVMs, cutaneous-mucosal venous malformations, and capillary malformations with AVM^71^. Specific signaling components are crucial for EC identity, including NOTCH, a key marker of arterial differentiation. Notably, the loss of signaling from either NOTCH or platelet-derived growth factor subunit B (PDGFB), has been associated with the formation of AVMs^72,73^ Additionally, SOX18 has been shown to regulate NOTCH1^74,75^, while the deletion of NR2F2 in HUVECs has been found to alter the expression of NOTCH2-4 and decrease PDGFB expression^15,73^. Collectively, these findings suggest that the NR2F2 and SOX18 signaling axis might be essential for regulating pathways implicated in vascular malformations. This study explores the potential of exploiting the NR2F2-SOX18 antagonistic relationship as a novel therapeutic target for treating vascular malformations.

The findings from this work establish a clear window of opportunity for therapeutic intervention in the context of congenital malformations acquired during fetal development. Vascular malformations present a unique challenge due to their complexity and location in the body, often rendering surgical intervention impractical^76^. In such cases, pharmacological therapies are the preferred option. An example is sirolimus (rapamycin), an mTOR inhibitor, which has shown positive results in some patients^77,78^. However, due to the heterogeneity of the disease, not all individuals respond to this therapy, highlighting the need for alternative pharmacological options^79^. This study suggests that blood vasculature plasticity during postnatal organogenesis, which relies on developmental pathways, can be leveraged to treat VUS. Here, we uncover that the balanced activity of SOX18 and NR2F2 is essential for the proper development of arteriovenous identity, providing new insights into the molecular mechanisms underlying vascular malformations and new pathways that may be exploited for future therapies.

In summary, this work offers a model of how paired molecular switches exert a balanced cooperative and competitive behaviour to instruct vascular development. By combining quantitative molecular imaging with patient-derived insights, we establish a framework for investigating other rare disorders in the context of transcription factors uncoupling.

## Methods and Materials

### Cell culture

Hela cells were cultured in supplemented DMEM (Gibco) with 10% FBS, 1% GlutaMAX (Gibco), 1% MEM Non-essential amino acids (MEM NEAA, Gibco). Cells were cultured at 37°C with 5% CO_2_. HUVEC cells were cultured in EBM-2 media (Lonza) supplemented with EGM-2 bullet kit (Lonza). Cells were cultured at 37 °C with 5% CO_2_.

### hESC

Undifferentiated H1-hESC (Wicell, WA01) were maintained in mTeSR™ plus medium (StemCell Technologies, #100-0276,100-1130) in monolayer culture on Geltrex (ThermoFischer Scientific, # A1413302). The medium was changed on alternate days per the manufacturer’s guidelines. The cells were split and maintained according to a previously published protocol^80^.

### Generation of H1-hESC with NR2F2^c994del^ mutation

H1-hESC containing NR2F2^c994del^ mutations were generated using CRISPR-Cas9 cells with synthetic gRNA (Synthego), Cas9 (Alt-R™ S.p. HiFi Cas9 Nuclease V3, IDT, #1081060) and HDR donor template (IDT). The knock-in strategy, gRNA sequence, and HDR oligo sequence are shown in **Supplementary** Figure 1C.

H1-hESCs at 40-50% were treated with 10 μM ROCK inhibitor (Thiazovivin; Tocris, #3845) 24h before the electroporation. Cells were then dissociated using Accutase, followed by dilution in DMEM/F12 (1:5) and centrifugation at 300g for 5 mins. Subsequently, cell pellets were resuspended in PBS and centrifuged at 300g for 5 mins. 200,000 cell pellets were then resuspended in 10 μl Buffer R (for each well in 24 wells plate). Cas9-gRNA complex was prepared by mixing 36μM Cas9 (final concentration) and 0.5 μl gRNA and incubated for 20 mins at RT. HDR donor oligo was added to the mix, followed by H1-hESC. Cells were electroporated using the NeonTransfection system (Fischer Scientific, Invitrogen™ #MPK5000) according to manufacturer guidelines (electroporation condition: 1400V, 20ms pulse width, 1 pulse). Following electroporation, cells were transferred to the Geltrex-coated 24-well plate containing 500 μl mTeSR plus + 10 μM Thiazovivin. The medium was changed to 24 hours post-transfection, followed by DMEM/F12 wash, and switched to mTeSR plus medium. Single cells were then FACS sorted (MA900), expanded, and Sanger sequenced to identify clones containing NR2F2^c994del^ mutation.

### Differentiation of H1-hESC to Artery and vein Endothelial cells

The protocol for differentiating artery and vein endothelial cells from H1-hESC has been published^42,80^. In short, at day 0, H1-hESCs were dissociated into single cells with Accutase (Sigma, A6964), and 175,000 cells were plated in each well of 12 wells of cell culture plates pre-treated with geltrex for 1h at 37°C. Cells were seeded in mTeSR plus + 1μM Thiazovivin (Tocris, #3845). On day 1, cells were washed briefly with DMEM/F12 (Thermo Fischer Scientific, 11320082) and differentiated in a primitive streak induction medium. On Day 2, cells were briefly washed and then differentiated in a lateral mesoderm induction medium (Day 2, lateral mesoderm). For Artery differentiation, cells were briefly washed and differentiated in an Artery induction medium for 24 h. For Veins, the Day 2 lateral mesoderm induction medium was incubated in day 3 vein intermediate medium, followed by day 4 vein medium.

| **Day 1: Mid primitive streak induction** | **Final** | **Stock** |
| --- | --- | --- |
| Activin A | 30 ng/mL | 50ug/ml |
| BMP4 | 40 ng/mL | 100ug/ml |
| CHIR99021 | 6 μM | 10mM (10mg/2.15ml) |
| FGF2 | 20 ng/mL | 100ug/ml |
| **Day 2: Lateral mesoderm** | **Final** | **Initial** |
| BMP4 | 40 ng/mL | 100ug/ml |
| GDC0941 | 2.5μM | 10mM |
| VEGF | 100 ng/mL | 100ug/ml |
| XA V939 | 1 μM | 10mM |
| Forskolin | 10 μM | 10mM |
| SB505124 | 2 μM | 10mM |
| AA2P | 200 μg/mL | 50mg/ml |
| **Day 3; ARTERY meidum** |  |  |
| Activin A | 15 ng/mL | 50ug/ml |
| GDC0941 | 2.5 μM | 10mM |
| VEGF | 100 ng/mL | 100ug/ml |
| XA V939 | 1 μM | 10mM |
| AA2P | 200 μg/mL | 50mg/ml |
| DMH1 | 250nM | 1mM |
| **Day 3,4: VEIN medium** |  |  |
| **Day 3 Vein medium** | **Final** | **Initial** |
| SB505124 | 2 μM | 10mM |
| DMH1 | 250nM | 1mM |
| RO4929097 | 2 μM | 10mM |
| VEGF | 100 ng/mL | 100ug/ml |
| XA V939 | 1 μM | 10mM |
| AA2P | 200 μg/mL | 50mg/ml |
| **Day 4 Vein medium** | **Final** | **Initial** |
| SB505124 | 2 μM | 10mM |
| RO4929097 | 2 μM | 10mM |
| PD0325901 | 0.5 μM | 5mM |
| CHIR99021 | 1 μM | 10mM |
| AA2P | 200 μg/mL | 50mg/ml |

### Rescue experiment of vein endothelial cells using R(+) Propranolol and Sm4

For rescue experiments, NR2F2^WT^ and NR2F2^c994del^ were treated with R(+) propranolol at 15 μM and Sm4 at 30 μM while differentiating cells in a day 4 vein medium, and cells were processed for FACS and qPCR analysis as described in the respective sections.

### Flow Cytometry

Cells were dissociated by incubation in Accutase for 5 mins at 37 ^°^C. Dissociated cells were diluted 1:5 in DMEM/F12+5% FBS (Thermo Fischer Scientific, # A5670401) and centrifuged at 350g for 5 minutes. Cell pellets were resuspended in FACS buffer (PBS+1mM EDTA+2% FBS+1% penicillin/Streptomycin). Cells were then blocked for 5 minutes in the presence of an FcR blocking reagent (Miltenyi Biotec, #130-059-901), followed by staining with PE anti-human CD73 (BioLegend, #344004) and FITC anti-human CD144 (BD Biosciences, #560411) for 30 minutes on ice, protected from light. Cells were then washed with FACS buffer (1:5), and resuspended in 300μl FACS buffer + DAPI (Miltenyi Biotec, # 130-111-570) for live/dead cell identification. Cells were then analyzed using a Sony MA900 Multi-application cell sorter. For analysis, cells were gated based on forward and side scatter, followed by height and width parameters to identify singlets. Finally, DAPI-negative (live cells) were gated for CD73 and CD144 populations.

### ATAC-seq

HUVECs were cultured in medium 199 (Sigma-Aldrich, MO, USA) supplemented with 16% fetal bovine serum (FBS), 100 U/ml penicillin, 100 U/ml streptomycin, 15 μg/ml endothelial cell growth factor (BD Biosciences, MA, USA) and 15 μg/ml heparin (Sigma-Aldrich, MO, USA) at a humidified 37 ◦C, 5% CO2 incubator.

HUVECs were grown to 60% confluence then treated with 25uM Sm4 or the equivalent volume of DMSO. 24 hours after treatment, 3 HUVEC lines were harvested and pooled. ATAC-seq and ATAC-seq analysis were performed by Active Motif. As per their sample preparation instructions, 150k cells/sample were collected, centrifuged at 500 x g at 4°C to remove the supernatant. Then, the cells were resuspended in ice-cold cryopreservation solution (50% FBS, 40% media, and 10% DMSO) and transferred to cryotubes for freezing. Samples were then shipped to Active Motif for analysis.

### HS17 sample preparation

The human embryo samples were collected for pathological examination, which entails histological verification of pregnancy and confirmation of fetal components. In addition, they were assessed to ensure the absence of conditions such as hydatidiform mole or choriocarcinoma. We selected residual specimens, collected between January 2000 and October 2021. The use of human samples was approved by the Ethics Committee of Mie University Hospital (approval number: H2021-228). Informed consent was acquired at the time of surgery, complemented by the availability of an opt-out provision, ensuring participants’ autonomy to withdraw from the study at any point. The staging of the embryos and fetuses used in the experiments was done using a combination of the Tissue collection and ethical considerations. For this study, 31 preserved human embryos in organogenesis period and 3 fetuses ranging from CS8 to GW9 were analyzed. The samples were collected for pathological examination, which entails histological verification of pregnancy (investigating the decidua’s interstitial and the endometrium’s Arias-Stella reactions) and confirmation of fetal components (examination of chorionic tissue). In addition, they were assessed to ensure the absence of conditions such as hydatidiform mole or choriocarcinoma. We selected residual specimens, collected between January 2000 and October 2021, that included embryos and utilized these specimens specifically for the purposes of this research.

### Immunohistochemistry (IHC)

#### Proband samples

Prior to staining the samples were processed to remove paraffin wax through multiple xylene and ethanol baths. Epitope retrieval was performed using Tris-EDTA. To stain the sample sections the samples were submerged in blocking buffer consisting of 1M maleic acid (Sigma, M0375), 10% heat inactivated horse serum (Sigma, H1138), 1% DMSO (Sigma, 472301), 0.1% PBTX (PBS + 0.1% Tween-20) (Tween-20, Sigma, P7949) for 1hr. Antibodies were then diluted into blocking buffer at 1:50 for anti-hCD31 (Abcam, ab28364) and 1:250 for anti-hNR2F2 (R&D systems, PPH714700) and anti-hSOX18 (Santa Cruz Biotechnology, sc-166025) overnight in the dark at 4C. The next day the slides were washed in blocking buffer three times over the course of 30min. Secondary antibodies were diluted in blocking buffer at 1:300 (ThermoFisher (Alexa Fluor 488 goat anti-rabbit IgG, A11008), (Alexa Fluor 555 goat anti-mouse IgG2a, A21137), (Alexa Fluor 647 goat anti-mouse IgG1, A21240)) for 1hr at room temp in the dark. Samples were then washed with PBS three times over 30min. Hoechst (ThermoFisher, R37605)) was diluted in PBS at 1:1000 and added to the slides for 10 min, then washed twice with PBS and mounted. Imaging was performed on a Zeiss 800 LSM. All images were processed using ImageJ software.

#### HS17 sample

IHC was performed using 3-μm thick sections. The sections were deparaffinised and rehydrated through a series of xylene and ethanol. To suppress autofluorescence, samples were incubated in 0.1% sodium borohydride in 0.1M phosphate-buffered saline (PBS) (137 mM NaCl, 2.7 mM KCl, 10 mM Na2HPO4, and 1.8 mM KH2PO4, pH 7.2) for 30 minutes, rinsed with water, and subsequently incubated for 5 minutes in 0.2M glycine in 0.1M PBS. Antigen retrieval was carried out using a pressure chamber with Tris-EDTA buffer (7.4 mM Tris, 1 mM EDTA-2Na, pH 9.0). Slides were incubated with primary antibodies against Prox1 (AF2727, R&D Systems, 1:150, RRID), Coup-TF2 (EPR18443, Abcam, 1:150, RRID), and SOX18 (sc-166025, Santa Cruz Biotechnology, 1:50, RRID). Alexa Fluor-conjugated secondary antibodies (Abcam, 1:400) were subsequently applied. Imaging was carried out using a Keyence BZ-X700 microscope. All images were processed using ImageJ software.

### Immunofluorescent staining of IH, VM and AVM FFPE patient tissue sections

FFPE tissue sections (5 μm) from patients with IH, VM or AVM were deparaffinized, immersed in an antigen retrieval solution (citrate-EDTA buffer containing 10 mM citric acid, 2 mM EDTA, and 0.05% Tween-20, pH 6.2) for 20 minutes at 95°C–99°C, cooled down and washed. Blocked for 30 min was performed in 10% donkey serum followed by incubation overnight at 4C with monoclonal IgG1 mouse anti-human SOX18 (D-8) (1:50, Santa Cruz Biotechnology, sc-166025), monoclonal IgG_2A_ mouse anti-human COUP-TF II (1:100, R&D, PP-H7147-00), and UEA1 fluorescently labeled with Alexa Fluor 649 (1:50, Vector Laboratories, DL1068). Next, the sections were incubated with Alexa Fluor 546 goat anti-mouse IgG1 (1:200; cross-absorbed, Invitrogen, A-21123) and Alexa Fluor 488 goat anti-mouse IgG_2A_ (1:200; cross-absorbed, Invitrogen, A-21137) as secondary antibodies for 45 minutes at room temperature. All slides were mounted using DAPI (Molecular Probes) to visualize nuclei. IF Images were acquired using a LSM 880 confocal microscope (Zeiss) through a 63x objective lens. All images were analyzed using Fiji ImageJ software.

### Immunofluorescent cell staining of HemSC, HemEC, and ECFC

Hemangioma stem cells (HemSC), hemangioma endothelial cells (HemEC) and endothelial colony forming cells (ECFC) were seeded on fibronectin-coated plates at a density of 30,000 cells/cm^2^ in EGM-2 media for 30 hours on 2 cm^2^ slides, fixed in 4% PFA and blocked in 5% BSA/0.3% Triton x-100 for 1 hour. Cells were co-stained for monoclonal IgG1 mouse anti-human SOX18 (D-8) (1:100, Santa Cruz Biotechnology, sc-166025), monoclonal IgG_2A_ mouse anti-human COUP-TF II (1:200, R&D, PP-H7147-00). Alexa Fluor 546 goat anti-mouse IgG1 (1:200; cross-absorbed, Invitrogen, A-21123) and Alexa Fluor 488 goat anti-mouse IgG_2A_ (1:200; cross-absorbed, Invitrogen, A-21137) served as secondary antibodies, incubation was for 45 minutes at room temperature. DAPI was used to visualize nuclei (Molecular Probes) followed by mounting (Invitrogen). Immunostainings of IH sections with respective isotype matched control IgG and secondary antibodies were conducted.

### RNA isolation and RT-qPCR

#### hESC

Total RNA was exracted using Trizol reagent (ThermoFisher Scientific, #1559608). RNA concentration was determined using Nanodrop (Thermoscientific). cDNA was produced using verso cDNA synthesis kit (ThermoFisher Scientific, 1453B). qPCR amplification was performed using PowerUP SYBR Green (ThermoFischer Scientific, A25778) on a Quantstudio 7 Flex (ThermoFischer Scientific). The 2^-ΔΔCT^ method was used to calculate relative gene expression relative to YWHAZ.

#### HUVEC drug treatment

The day before collection 300k cells/well were seeded in a 6 well plate. The following day HUVECs were treated with either CIA1 (Tocris, 7494) at 4μM or CIA1 at 4μM + R(+)propranolol (Sigma, P0689) at 25μM for 4 hours prior to collection. After 4 hours cells were washed with PBS. RNA was extracted using Trizol reagent (ThermoFisher, #1559618). RNA concentration was determined using Nanodrop (Thermo Scientific). cDNA was produced using cDNA kit (Bio-Rad, 1725035) and qPCR amplification was performed using SensiFAST SYBR kit (Bio-Rad, BIO-98050) on a Roche Lightcycler 480 II. The 2^-ΔΔCT^ method was used to calculate relative gene expression relative to GAPDH.

#### IH cells

Total RNA was extracted from cells with the RNeasy Micro Extraction Kit (QIAGEN). Reverse transcriptase reactions were performed using an iScript cDNA Synthesis Kit (Bio-Rad). qPCR was performed using SYBR FAST ABI Prism 2× qPCR Master Mix (Kapa BioSystems). Amplification was carried out in a QuantStudio 6 Flex Real-Time PCR System (Fisher Scientific). A relative standard curve for each gene amplification was generated to determine the amplification efficiency, with greater than 90% considered acceptable. Fold increases in gene expression were calculated according to the ΔΔCt method, with each amplification reaction performed in duplicate or triplicate. ATP5B was used as housekeeping gene expression reference. A list of all primer sequences used in this study is attached below.

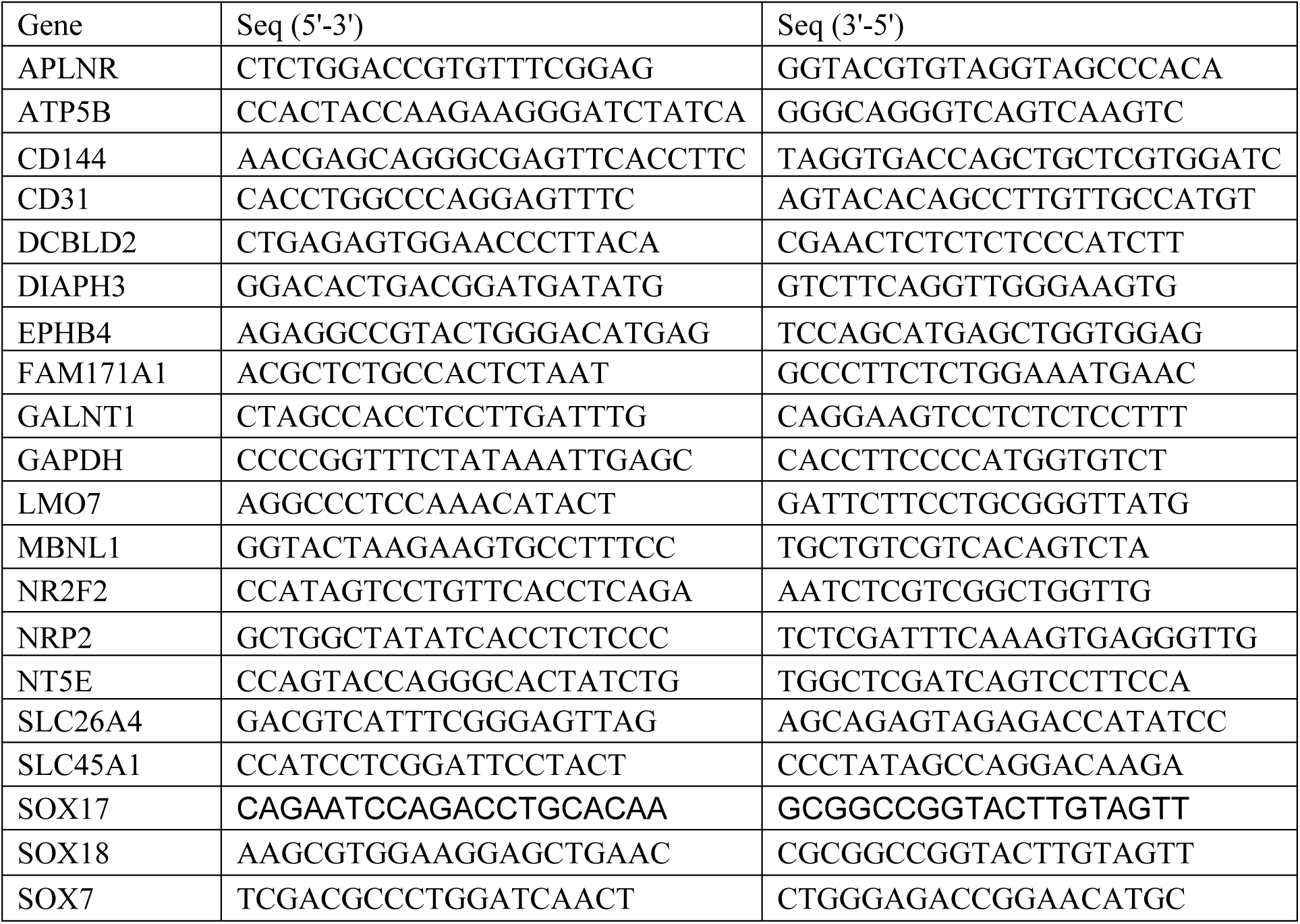
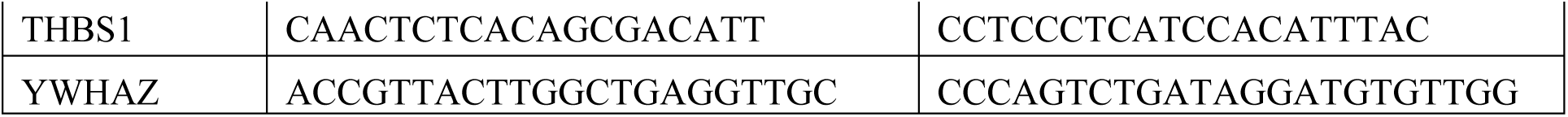

#### Primers

##### NR2F2^c994del^ Plasmid creation

Halo-NR2F2^WT^ (Genecopoeia, EX-C0221-M49) was used as a template to construct the c994del variant plasmid using the megaprimer sight-directed mutagenesis method. The Halo-NR2F2^c994del^ plasmid was validated by sanger sequencing to ensure the deletion occurred at position at 994.

### Quantitative molecular imaging Single-molecule tracking (SMT)

#### SMT Sample preparation

Single-molecule tracking was performed as described in McCann et al.^48^. HeLa cells were seeded at a density of 23,000 cells/well in 8-well chamber glass slides (Ibidi, 80827) coated with 0.5% gelatine 24h prior to transfection. 300ng of plasmid DNA/well of either Halo-tagged NR2F2^WT^ or NR2F2^c994del^ was transiently transfected into the cells using X-tremeGENE 9 Transfection Reagent kit (Roche, XTG9RO). For dual transfection experiments 600ng (300ng of each plasmid) of plasmid DNA/well of Halo-tagged NR2F2^WT^ + eGFP-SOX18, NR2F2^c994del^ + eGFP-SOX18, Halo-tagged SOX18 + eGFP-NR2F2^WT^, or SOX18 + eGFP-NR2F2^c994del^. Cells were then incubated at 37°C with 5% CO_2_ overnight prior to imaging then washed three times and then incubated in imaging media (Fluorobrite DMEM (Glibco), 10% FBS, 1% HEPES, 1% GlutaMax) 45 min prior to imaging 1nM of JF549 Halo-tag ligand (Promega, GA1111) was added directly to the media and cells were incubated for 10 minutes at 37 ° C with 5% of CO_2_. Following incubation, cells were washed twice 15min apart and replaced with imaging media.

#### SMT acquisition

Images were acquired on a Nikon TIRF microscope at a TIRF angle of 61 degrees to achieve HILO illumination. Samples were recorded with an iXon Ultra 888 EMCCD camera, filter cube TRF49909 – ET – 561 laser bandpass filter and 100 X oil 1.49 NA TIRF objective. Cells were imaged using a 561nm excitation laser at a power density of 10.3 μW to perform two different acquisition techniques. A fast frame rate which uses a 50 Hz (20ms acquisition speed) to acquire 6000 frames without intervals to measure displacement distribution and fraction bound, and a slow frame rate which uses a 2 Hz (500ms acquisition speed) to acquire 500 frames without intervals to measure residence times.

#### SMT analysis

Masking and segmentation of the nucleus was performed in ImageJ for all files. To identify and track molecules a custom-written MATLAB implementation of the multiple target tracing (MTT) algorithm, known as SLIMfast was used^81^. Parameters used for fast frame rate analysis: Localization error: 10^-6.5^, blinking (frames) = 1, max # of competitors: 3, max expected diffusion coefficient = 3μm^2^/sec, box size = 7, timepoints = 7, clip factor = 4. Cells with less than 500 trajectories based on the above parameters were excluded from analysis. The first four frames of each trajectory were used to calculate the mean squared displacement. Diffusion coefficient was calculated from each trajectory’s mean squared displacement and plotted. An inflection point was determined on WT NR2F2 diffusion profile and used as a boundary to determine confined vs non-confined states for all conditions. The confined fraction and non-confined fraction of each cell were calculated by computing the number of trajectories whose mean diffusion coefficient were defined as confined and non-confined respectively. Parameters used for slow frame rate analysis: Localization error: 10^-7^, blinking (frames): 1, max # of competitors: 3, max expected diffusion coefficient = 0.33μm^2^/sec, box size = 9. Slow-tracking analysis was performed using custom MATLAB code based on ^52^.

### Fluorescence fluctuation spectroscopy (FFS)

#### Halo Tag labelling for FFS experiment

To saturate all Halo Tags with Janelia Farm dyes 549 or 646 for single or dual-color labelling, HeLa cells transiently expressing Halo Tag-tagged Sox7, NR2F2 or NR2F2_C994del constructs were incubated with a saturating concentration of a single JF dye (JF549, 100 nM) or two JF dyes (JF549 : JF646, 100 nM : 100 nM) for 15 minutes at 37 °C, similar to McCann et al.^48^. Cells were washed twice with 1× PBS before imaging experiments.

#### FFS acquisition

All FFS measurements for Number and Brightness (NB) analysis and cross Raster Image Correlation Spectroscopy (RICS) were performed on an Olympus FV3000 laser scanning microscope coupled to an ISS A320 Fast FLIM box for fluorescence fluctuation data acquisition. For single channel NB FFS measurements, HaloJF549 tagged plasmids were excited by a solid-state laser diode operating at 561 nm and the resulting fluorescence signal was directed through a 405/488/561 dichroic mirror to an external photomultiplier detector (H7422P-40 of Hamamatsu) fitted with a mCherry 600-640 nm bandwidth filter. For dual channel RICS FFS measurements (that enable cross RICS), the HaloJF546 and HaloJF646 combination were excited by solid-state laser diodes operating at 561 nm and 640 nm, respectively, and the resulting signal was directed through a 405/488/561/640 dichroic mirror to two internal GaAsP photomultiplier detectors set to collect 600-640 nm and 650-750 nm, respectively.

All FFS data acquisitions (i.e., NB, cross RICS) employed a 60X water immersion objective (1.2 NA) and first involved selecting a 10.6 μm region of interest (ROI) within a HeLa cell nucleus at 37 °C in 5% CO2 that exhibited low protein expression level (nanomolar) to ensure observation of fluctuations in fluorescence intensity^82^. Then a single or simultaneous two channel frame scan acquisition was acquired (N = 100 frames) in the selected ROI with a pixel frame size of 256 x 256 (i.e., pixel size ∼ 41nm) and a pixel dwell time of 12.5 µs. These conditions resulted in scanning pattern that was found to be optimal for simultaneous capture of the apparent brightness and mobility of the Halo tagged constructs being characterised by NB and cross RICS analysis; all of which was performed in the SimFCS software developed at the Laboratory for Fluorescence Dynamics (LFD).

#### Number and brightness (NB) analysis

The oligomeric state of the different HaloJF549-tagged plasmids investigated was extracted and spatially mapped throughout single channel FFS measurements via a moment-based brightness analysis that has been described in previously published papers ^83,84^. In brief, within each pixel of an NB FFS measurement there is an intensity fluctuation F(t) which has: (1) an average intensity 〈F(t)〉 (first moment) and (2) variance *σ*^2^ (second moment); and the ratio of these two properties describes the apparent brightness (B) of the molecules that give rise to the intensity fluctuation. The true molecular brightness (*ɛ*) of the fluorescent molecules being measured is related to B by B = ε + 1, where 1 is the brightness contribution of a photon counting detector. Thus, if we measure the B of monomeric HaloJF549-Sox7 (B_monomer_= *ɛ_monomer_* + 1) under our NB FFS measurement conditions, then we can determine *ɛ_monomer_* and extrapolate the expected B of HaloJF549-tagged dimers (B_dimer_= *(2 x ɛ_monomer_)* + 1) or oligomers (e.g., B_tetramer_= *(4 x ɛ_monomer_) + 1), and in turn define* brightness cursors, to extract and spatially map the fraction of pixels within a NB FFS measurement that contain these different species. These cursors were used to extract the fraction of HaloJF549-NR2F2 dimer and oligomer (i.e., number of pixels assigned B_dimer_ or B_oligomer_) within a NB FFS measurement and quantify the degree of NR2F2 self-association across multiple cells. Artefact due to cell movement or photobleaching were subtracted from acquired intensity fluctuations via use of a moving average algorithm and all brightness analysis was carried out in SimFCS from the Laboratory for Fluorescence Dynamics.

#### Cross RICS analysis

The fraction of interaction between the HaloJF549 and HaloJF646-tagged NR2F2 or NR2F2-C994del plasmids was extracted via application of cross RICS functions described in previously published papers to the dual channel FFS measurements^48,82^. In brief, the fluorescence intensity recorded within each frame (N =100) of each channel (i.e., CH1 and CH2) was spatially correlated via application of the RICS function, and spatially cross-correlated between channels (CC) via application of the cross RICS function, alongside a moving average algorithm (N = 10 frames) in both instances. Then the recovered RICS and cross RICS correlation profiles were fit to a 3D diffusion model and the amplitude versus decay of each fit recorded in the form of a G value and diffusion coefficient (D) respectively. The ratio of the cross RICS amplitude (i.e., G_CC_) with the limiting channel RICS amplitude (i.e., G_CH1_ or G_CH2_) enabled the fraction of HaloJF549-NR2F2 molecules interacting with Halo646-NR2F2 molecules to be extracted. All cross RICS analysis was carried out in SimFCS from the Laboratory for Fluorescence Dynamics.

### Statistics

All statistical analyses were performed using GraphPad Prism. An outlier test using settings (Q=1) was performed to identify and remove outliers. Results are displayed as mean ± the standard deviation unless indicated otherwise.

### Study approvals and ethics

#### Proband samples

The use of the proband’s samples was approved by the Hunter New England Exemption from HREC case study (authorization number: AU2206-11). All procedures were conducted in compliance with guidelines and regulations. ***hESCs*** The use of human embryonic stem cells (hESCs) in this study was approved by Stem Cell Research Oversight (SCRO) at Stanford University. All procedures were conducted in compliance with institutional guidelines and regulations. ***CS17 sample*** The use of human samples was approved by the Ethics Committee of Mie University Hospital (approval number: H2021-228). Informed consent was acquired at the time of surgery, complemented by the availability of an opt-out provision, ensuring participants’ autonomy to withdraw from the study at any point. ***HemSC, HemEC, and ECFC*** Vascular anomaly specimens were obtained under protocols approved by the Committee on Clinical Investigation at Boston Children’s Hospital (IRB protocol number 04-12-175R and P00003505; AH, JB). Patient specimens were collected upon written informed consent of the patient or the patient’s guardian, de-identified, and used for cell isolation as well as tissue staining under a Boston Children’s Hospital IRB approved protocol (04-12-175R and P00003505) and in accordance with Declaration of Helsinki principles. ***HUVEC cells*** Umbilical cords were collected from the Royal Prince Alfred Hospital upon written informed consent of the patient or the patient’s guardian, de-identified, and used for cell isolation of human umbilical vein endothelial cells (HUVECs). Isolation protocol was as previously described in Litwin et al.^85^ and cells were used from passage 1 to 3. Ethics approval for the collection of the umbilical cords was from SLHD Protocol No. X16-0225.

## Data Availability

The data that supports the findings in this study are available at DOI: 10.5281/zenodo.15098820.

## One-sentence summary

Antagonistic interplay between SOX18 and NR2F2 allows therapeutic rescue of a rare NR2F2 mutation syndrome using a SOX18 inhibitor.

## Acknowledgments

We thank the Vascular Anomalies Center and Department of Pathology at Boston’s Children’s Hospital for patient specimens. We acknowledge the technical and scientific staff of Sydney Microscopy & Microanalysis, the University of Sydney node of Microscopy Australia. Research reported in this manuscript was supported by NIH-R01HL12850307 (SJ), University of Sydney Drug Discovery Initiative - seed funding (grant ID: 23059) (MG, PC), Australian Excellence in Diversity Fellowship (JQ), the Vascular Anomalies Center at Boston Children’s Hospital (AH), and the Grants-in-Aid for Scientific Research from the Ministry of Education, Culture, Sports, Science, and Technology of Japan (23K15949) and Japan Agency for Medical Research and Development (AMED) (22jm0610079h0001) (KM). KRH was supported by NIH-R01HL12850307 and the Howard Hughes Medical Institute. NHMRC Idea Grants (APP2019904 and APP2029719) (MF)

## Author contributions

MG, SJ, MF, and KRH designed the study. MF coordinated the studies with all laboratories and provided the original idea of pharmacological rescue. GM coordinated the interaction between SK and MF. MG, SJ, JQ, AH, PC, TK, WL, and KM conducted experiments. MG, SJ, JQ, AH, YW, and KM carried out formal analysis. AH, SN, ES, TD, and KM provided samples, clinical management, and ethics. MG, AH, EH, and KM performed data curation. MG and MF wrote the original manuscript. All authors reviewed, edited, and agreed to the final version of the manuscript. MF and KRH acquired funding.

---

**Supp Fig 1:**
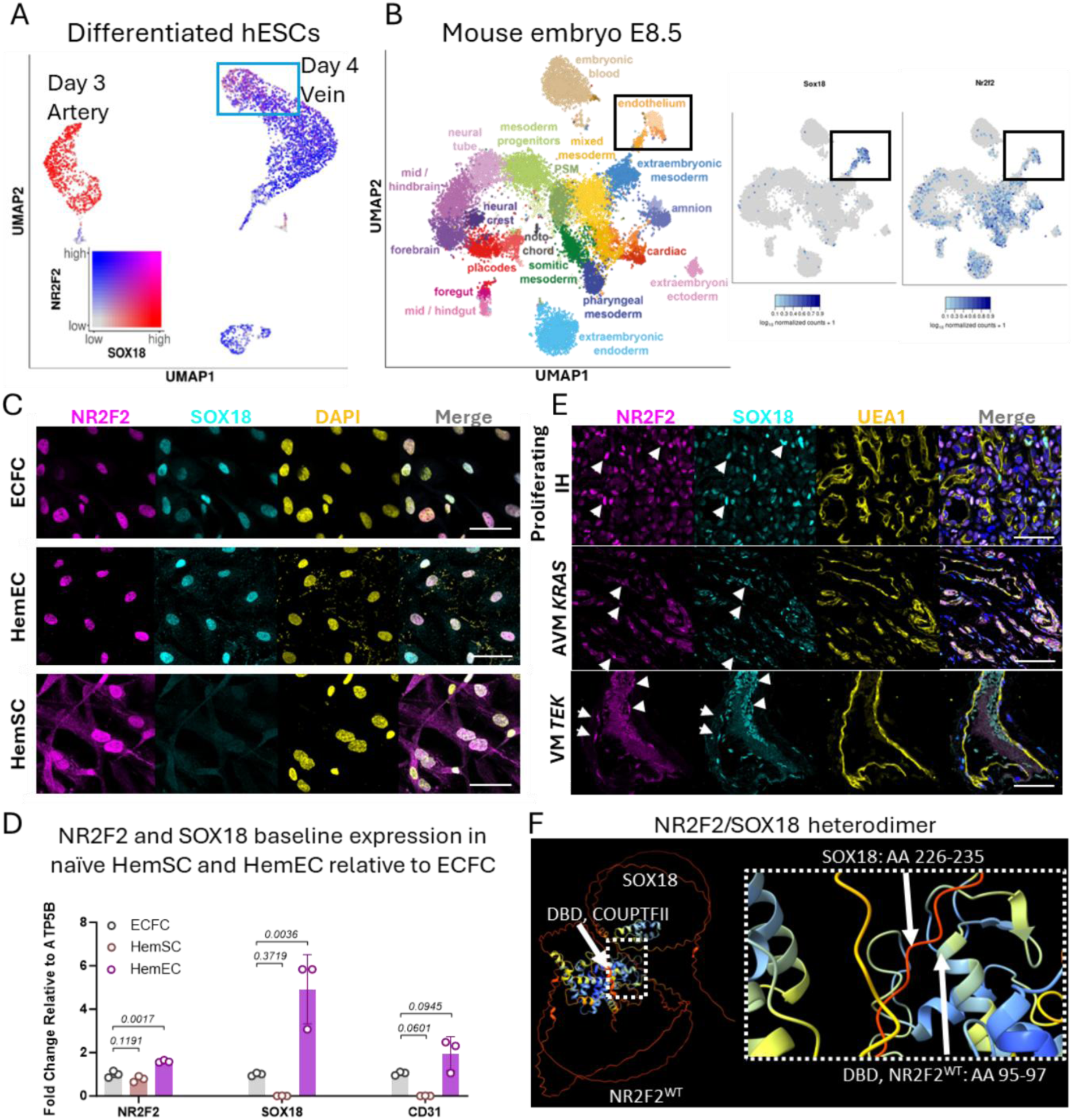
*In vivo* SOX18 and NR2F2 are co expressed during human development and vascular malformations. **A** UMAP of NR2F2 and SOX18 from scRNA-seq dataset from Ang et al. of ECs derived from hESCs. Blue box denotes venous population expressing both NR2F2 and SOX18. **B** UMAP of NR2F2 and SOX18 from scRNA-seq dataset from Ibarra-Soria et al. of mouse embryo at E8.5. **C** Immunofluorescent staining of NR2F2 (magenta) and SOX18 (cyan), and nuclei stained with DAPI (yellow) in IH patient-derived hemangioma stem cells (HemSC), hemangioma endothelial cells (HemEC), and ECFC demonstrates nuclear co-expression (scale bars 25 µm). **D** qPCR comparing mRNA levels of NR2F2 and SOX18 in patient-derived HemSC and HemEC relative to ECFC (n=3 biological replicates for HemSC and HemEC, n=3 independent experiments for ECFC; P values were calculated using one-way ANOVA with Dunnett’s multiple comparisons test. Data show the mean ± SD. **E** Immunofluorescent staining of NR2F2 (magenta), SOX18 (cyan), and the endothelial marker UEA1 (yellow) in IH, VM, and AVM patient tissue. White arrows denote nuclear co-expression of NR2F2 and SOX18 in EC (scale bars 50 µm). **F** Alphafold 2 modeling of NR2F2^WT^ and SOX18 (left). Dimer formation occurs at amino acids 95-97 (DNA binding domain) on NR2F2^WT^ and at amino acid 226-235 (TAD) on SOX18.

**Supp Fig 2:**
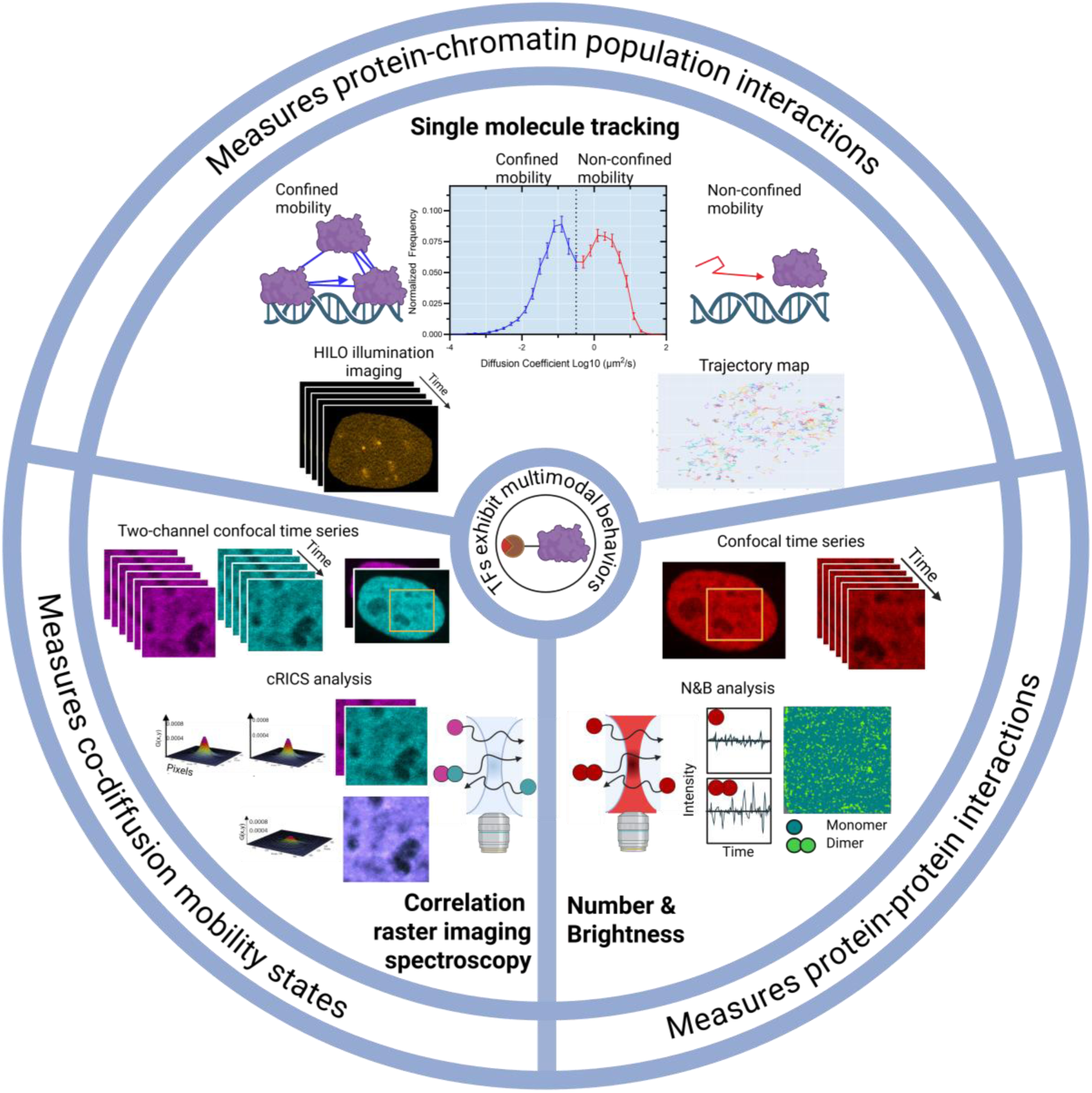
Biophysical features of transcription factors behavior involved in gene regulation. TFs exhibit distinct multimodal behaviors that reflect their functional roles, including chromatin interactions and PPI. To investigate these behaviors we applied the same imaging approaches described in McCann et al. Aligning the various dynamic behaviors provides a more comprehensive understanding of how the TFs navigate and interact within the nuclear environment. **Single molecule tracking (SMT)**. SMT was performed by acquiring time-lapse images of sparsely labeled fluorescent molecules using HILO illumination. In each frame, the position of individual fluorescent molecules was identified, and trajectories were reconstructed by linking positions across consecutive frames. The trajectories provide a dynamic map of molecular motion from which mean squared displacement and diffusion coefficients are calculated. To distinguish different mobility states, diffusion coefficients are plotted as a distribution and a threshold is set to at the inflection point between two peaks representing confined (low diffusion, blue) and non-confined (high diffusion, pink) motion. The proportion of molecules in each mobility state is then calculated to determine changes in nuclear dynamics across conditions. **Number and brightness (N&B).** N&B analysis is conducted using confocal microscopy to acquire a time series of images of saturated labelled fluorescent molecules. The oligomeric state of the TFs is inferred by analysing fluctuations in fluorescence intensity at each pixel through time. These fluctuations are used to calculate brightness values, which scale with the number of fluorescent molecules, in such a way that a dimer is approximately twice as bright as a monomer. Monomer calibration was assessed using a TF known to exist only in a monomeric state, allowing extrapolation to identify dimer and higher order oligomers. **Cross-raster image correlation spectroscopy (cRICS).** Similar to N&B, cRICS involves acquiring a time series of images of saturated fluorescently labelled molecules. However, cRICS simultaneously captures two channels (different fluorophores) at high spatial resolution, ensuring that oversampling of the point spread function occurs. The RICS function computes spatial correlations of fluorescence fluctuations within each individual channel, revealing the diffusion behaviour of molecules in that channel. The cRICS function extends this by cross-correlating intensity fluctuations between the two channels, allowing detection of co-diffusing molecules. By comparing the diffusion coefficients obtained from the RICS (single channel) and cRICS (two channel), its possible to infer whether proteins are diffusing as monomers or as part of a complex, enabling quantification of interaction dynamics.

**Supp Fig 3:**
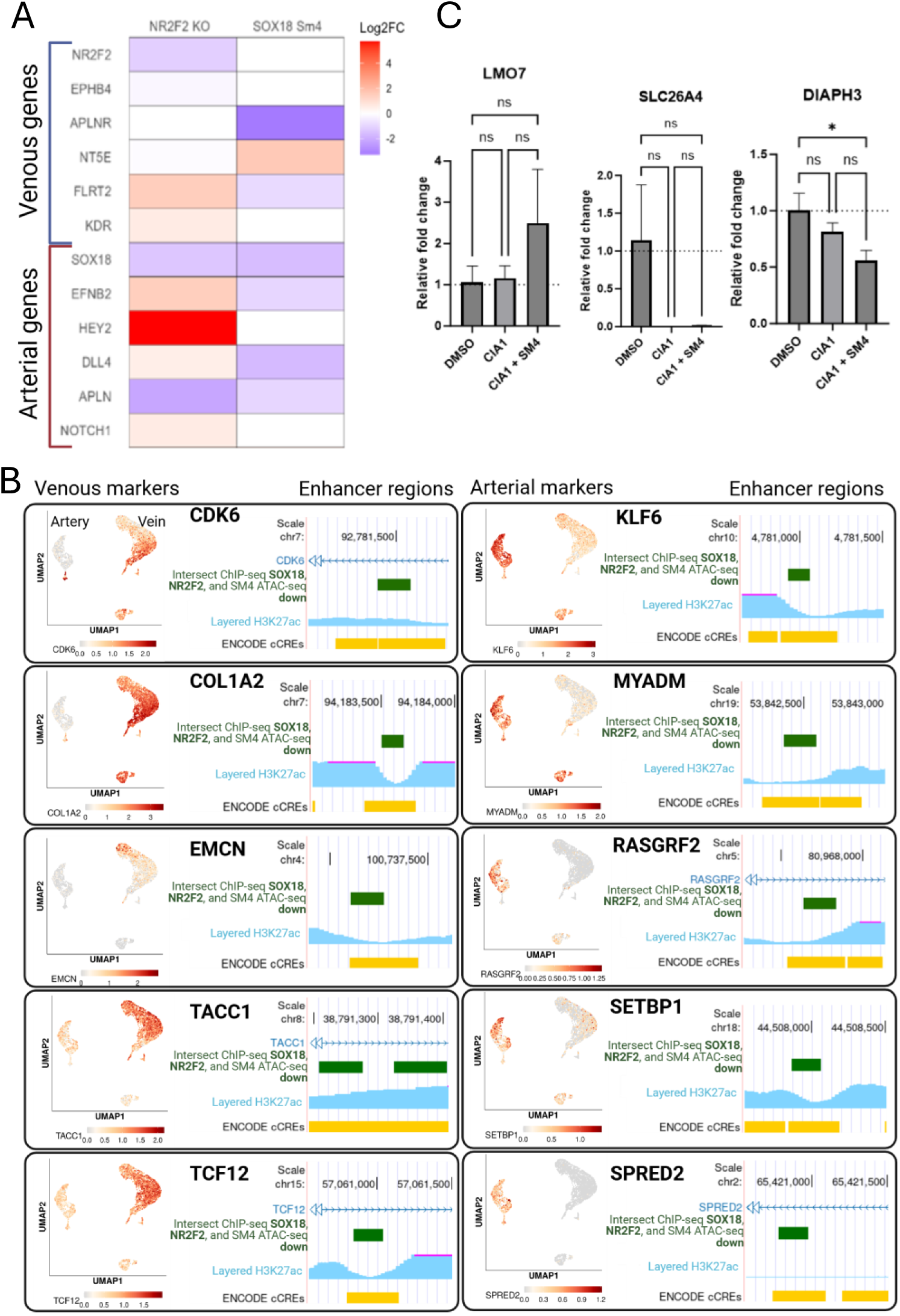
NR2F2 and SOX18 co-regulate a set of genes that determine endothelial arterio-venous identity. **A** Expression level of a subset of genes commonly used to distinguish arterial and venous endothelial cells identity. Datasets are from publicly available scRNA-seq datasets (NR2F2 knockout (KO), Sissaoui et al., and SM4 treatment of SOX18, (Overman et al.) **B** Subset of genes that are identified as either venous or arterial based on scRNA-seq dataset (Ang et al.) correlated with two publicly available ChIP-seq datasets for either SOX18 (McCann et al.) and NR2F2 (Sissaoui et al.) intersected with an ATAC-seq dataset of HUVECs treated with SM4. **C** Relative gene expression of treated HUVECS measured by qPCR. For panel C (n = 3 repeats), statistical analysis was performed using one-way ANOVA, and significance between groups was determined using Tukey’s multiple comparison post hoc test, * p < 0.05.

**Supp Fig 4:**
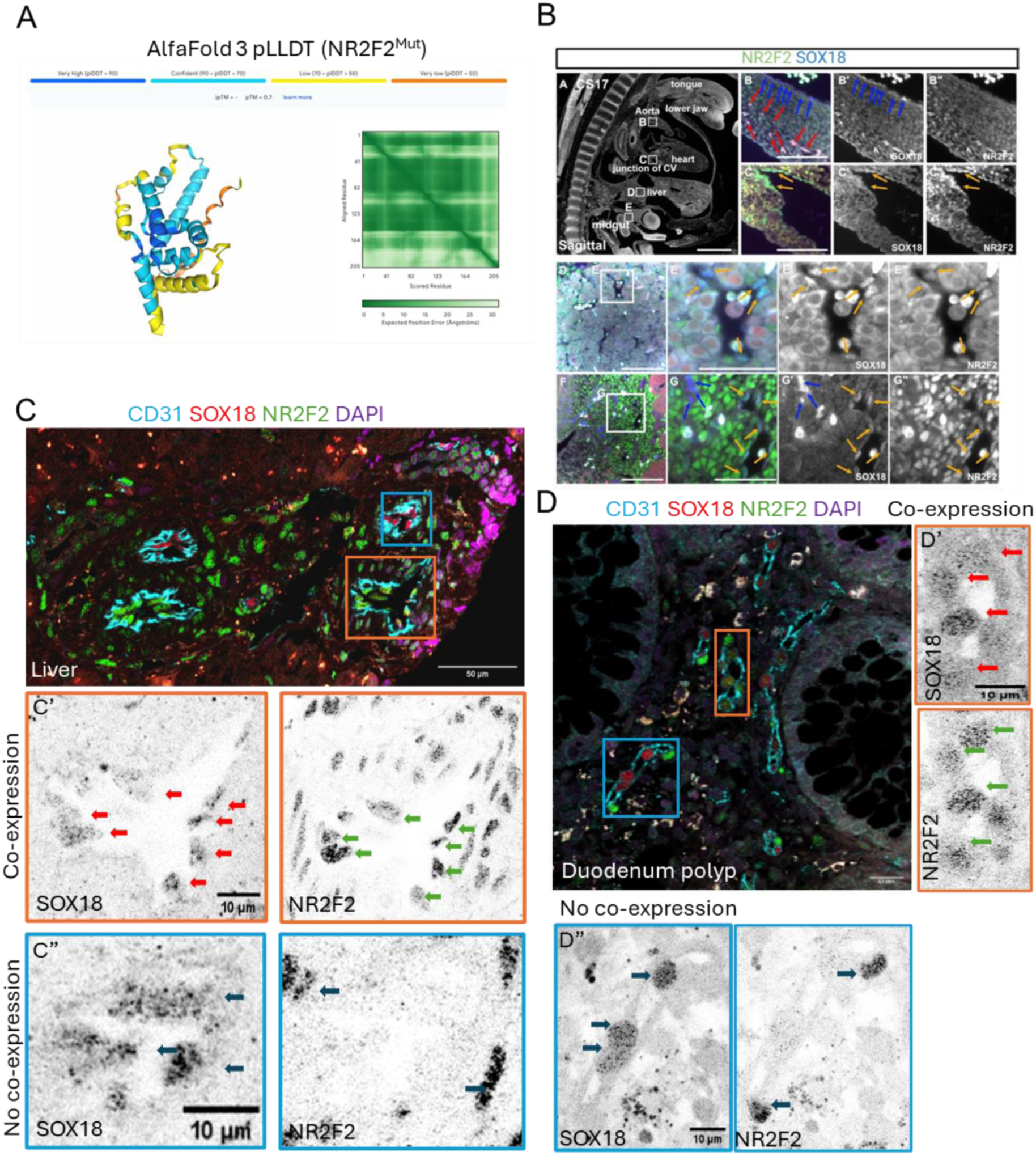
NR2F2 and SOX18 are co-expressed in human ECs during embryogenesis and in the proband tissues. **A** The predicted local distance difference tests (pLDDT) for the LBD of NR2F2^Mut^. **B** Immunostaining of NR2F2 (green) and SOX18 (blue) in sagittal sections from a CS17 embryo (A-D”’). The immunostaining shows co-expression of NR2F2 and SOX18 in endothelial cells during human development. **C-C**” and **D-D**” Sections of proband liver (C) and duodenum (D) stained for CD31 (cyan), SOX18 (red arrows), NR2F2 (green arrows), and DAPI (purple). Orange and blue boxes denote zoomed in regions of individual vessels. Orange box represents vessel that shows cells co-expressing both NR2F2 and SOX18 in the same nucleus, whereas the gold box is a vessel with cells either expressing NR2F2 or SOX18. Scale bar = 50μm (C-D) and 10 μm (C’”-D’”).

**Supp Fig 5:**
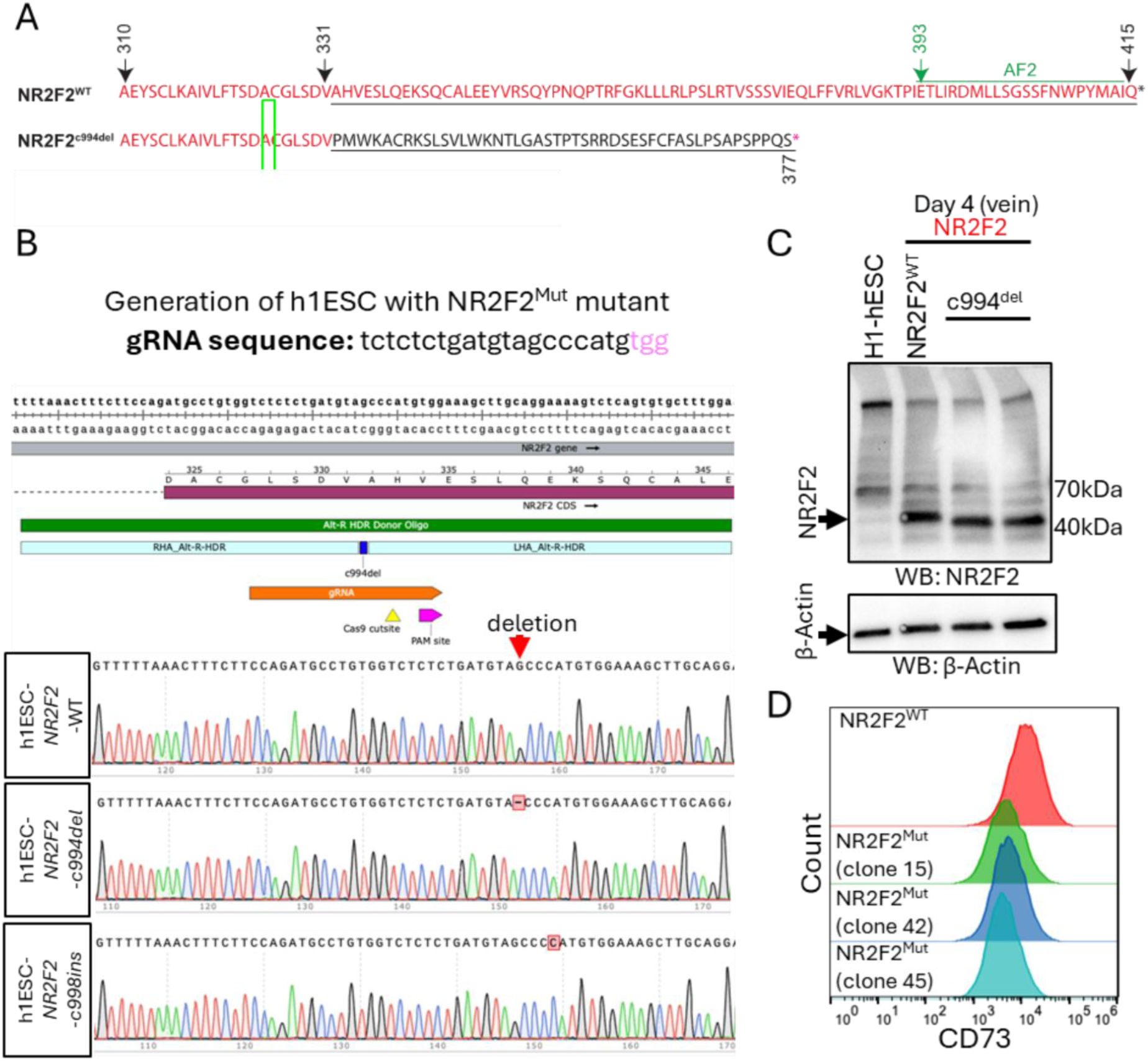
Generation of hESC with NR2F2^Mut^ variant. **A** Amino acid sequence comparison of NR2F2^WT^ and NR2F2^Mut^, NR2F2^c998ins^, an unrelated CRISPR modified cells used as control. **B** Generation of H1-hESC mutant clones harboring the NR2F2^Mut^ variant and an unrelated clone NR2F2^c998ins^. **C** Western blot showing NR2F2^Mut^ protein expression levels compared to wild type in hESC derived venous cells. **D** Histogram plots of CD73 expression on day four venous endothelial cells derived from different CRISPR-clones of NR2F2^Mut^ h1ESCs.

**Supp Fig 6:**
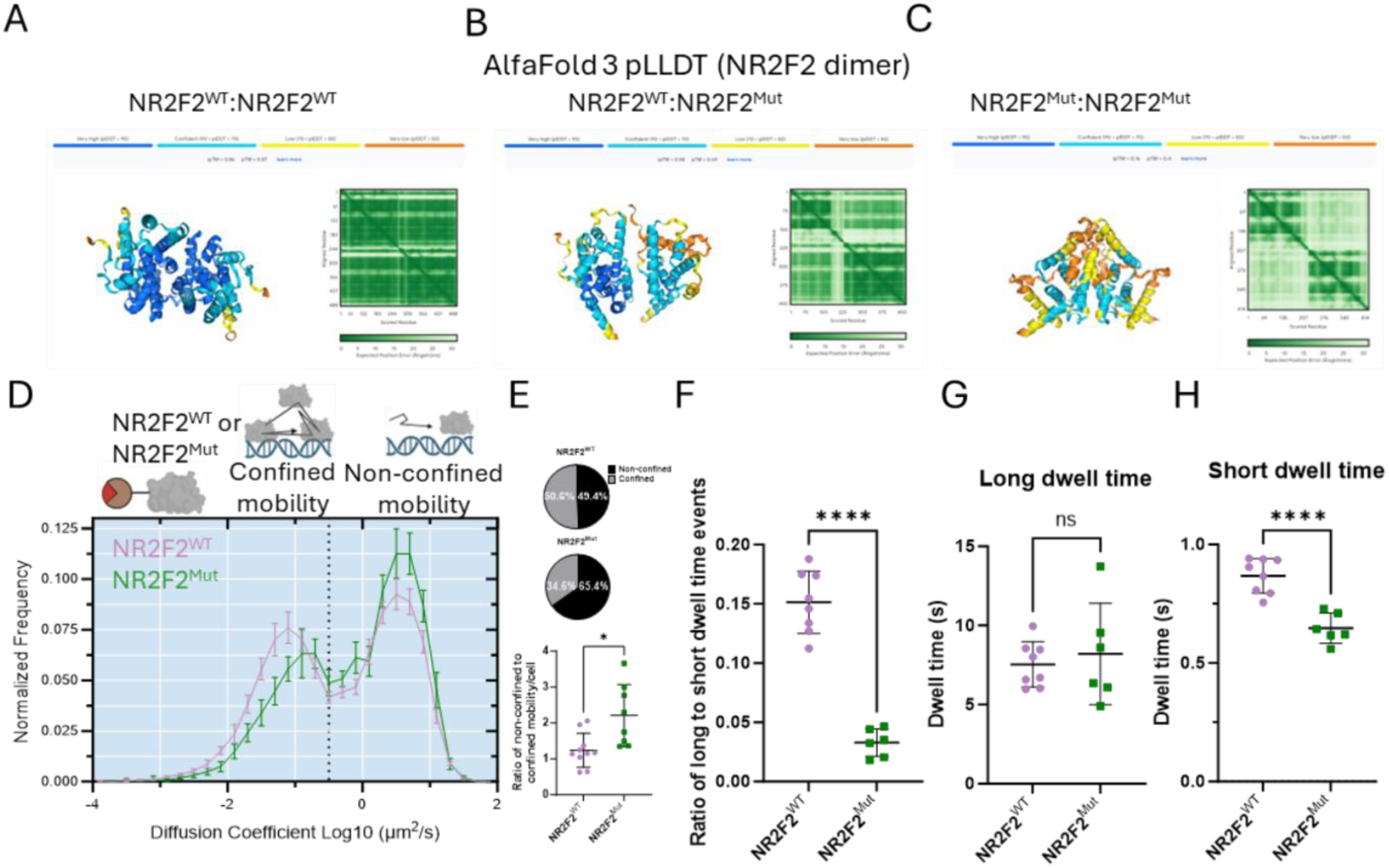
*In silico* modeling of NR2F2 LBD of variant shows loss of interaction. **A**-**C** The predicted local distance difference tests (pLDDT) for dimerization of LBD NR2F2^WT^ (A) and dimerization of NR2F2^WT^, NR2F2^Mut^ (B) and dimerization of NR2F2^Mut^ (note low confidence in the structure). **D** Diffusion mobility graph from SMT comparing NR2F2^WT^ and NR2F2^Mut^ in HUVECs. **E** Pie charts represent the proportion of the population that falls into either confined mobility or non-confined mobility based on diffusion coefficient. Ratio of non-confined to confined molecules per cell from C. **F-H**) Temporal occupancy characteristics. E) Ratio of long to short dwell times, F) long dwell time, and G) short dwell times. For panels D-G, n > 6 cells, statistical significance was determined by Welch’s T-test, * p < 0.05, **** p < 0.0001.

**Supp Fig 7:**
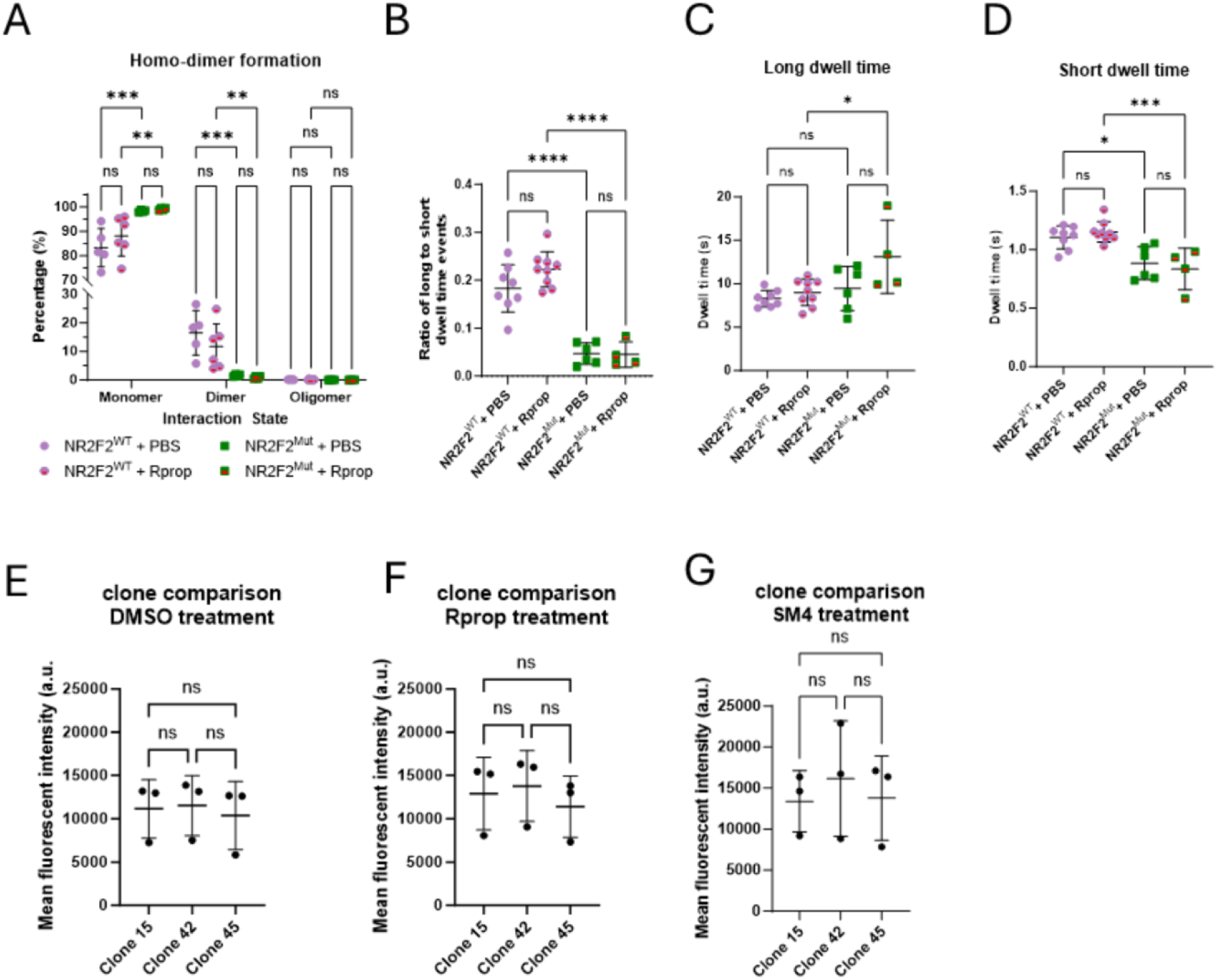
Dimer formation and temporal occupancy are not affected by propranolol treatment and hESC NR2F2^Mut^ clones react in a similar way to SOX18 inhibition. **A** Monomer, dimer, and oligomeric states measured by N&B. **B-D** Temporal occupancy characteristics. D Ratio of long to short dwell times, E long dwell time, and F short dwell times. **E-G** CD73 expression on NR2F2^Mut^ cells treated with either R(+)propranolol or SOX18. Statistical analysis was performed using a two-way ANOVA, statistical significance between groups was determined using Tukey’s multiple comparison post hoc test (A), a one-way ANOVA (B-D), statistical significance between groups was determined using Tukey’s multiple comparison post hoc test and a Kruskal-Wallis test (E-G), statistical significance between groups was determined using Dunn’s multiple comparison test , * p < 0.05, ** p < 0.005, *** p < 0.0005, and **** p < 0.0001.

**Supp Table 1:**
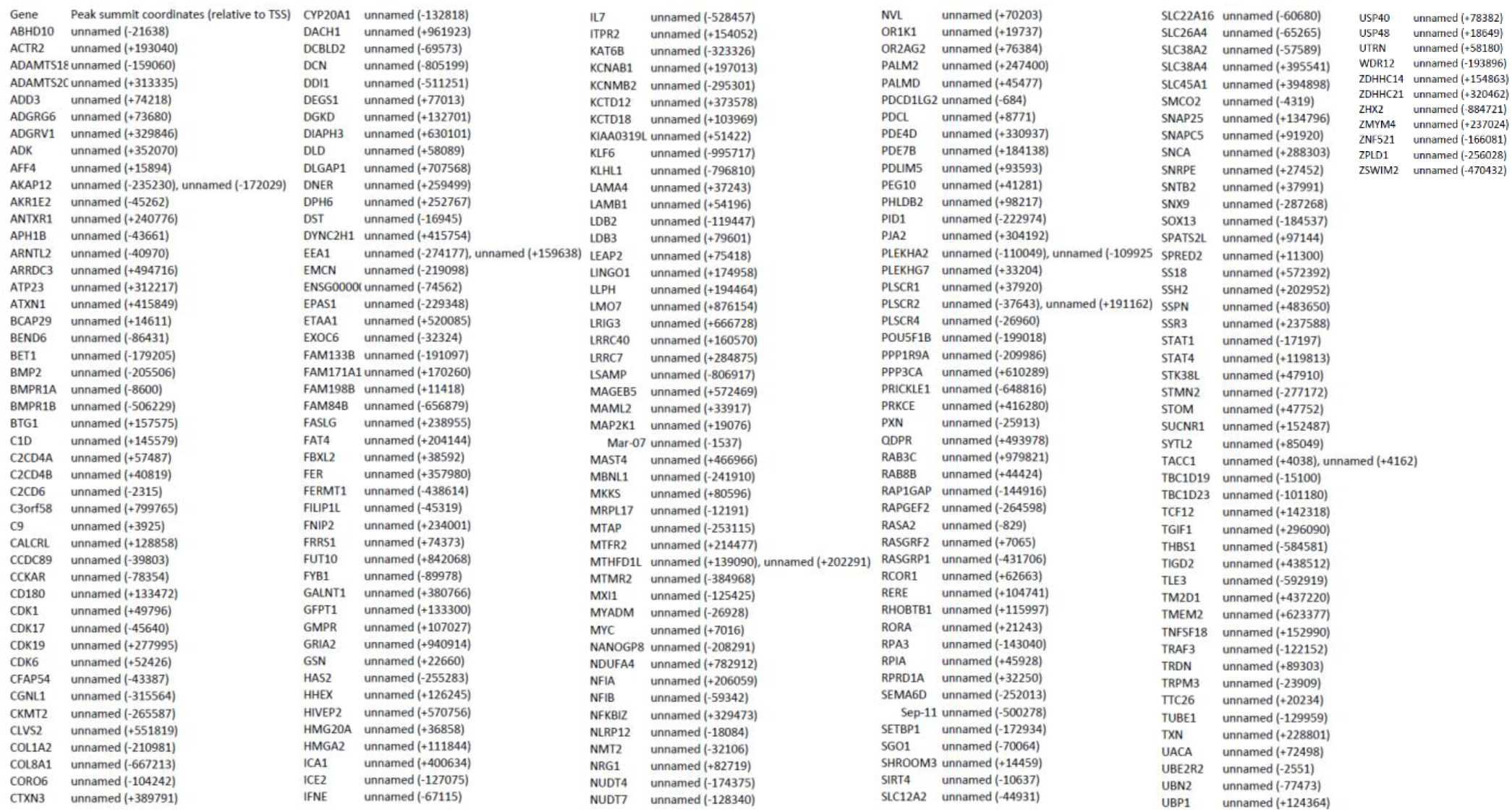
Peak to gene analysis of assigned enhancers relative to TSS from intersected peaks with ATAC-seq and ChIP-seq datasets.

**Supp Table 2:**
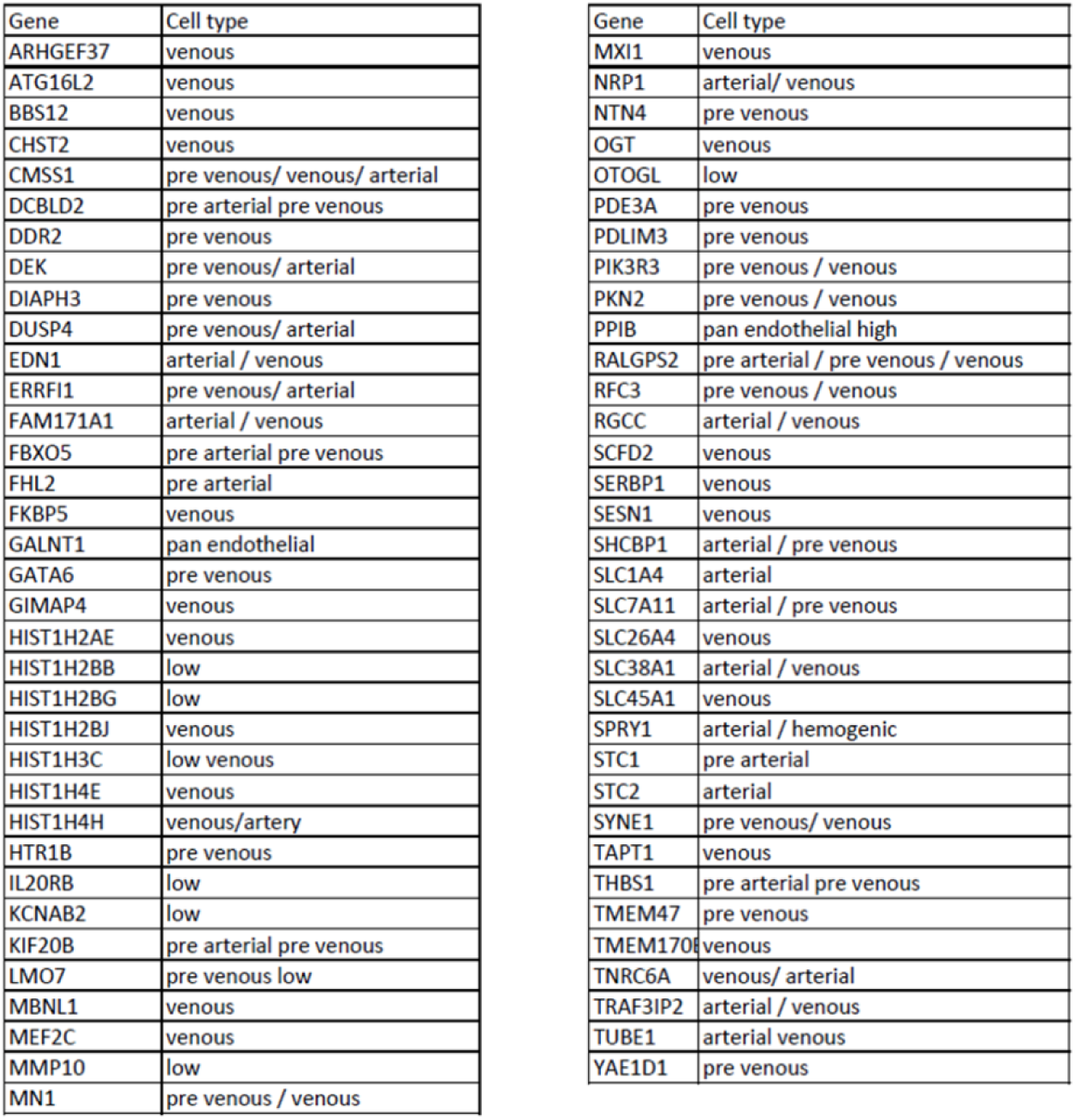
Subset of genes from Supp Table 1 that have arterial or venous specific profiles.

